# Arid5a uses disordered extensions of its core ARID domain for distinct DNA- and RNA-recognition and gene regulation

**DOI:** 10.1101/2024.02.29.582703

**Authors:** Julian von Ehr, Lasse Oberstrass, Ege Yazgan, Lara Ina Schnaubelt, Nicole Blümel, Francois McNicoll, Julia E. Weigand, Kathi Zarnack, Michaela Müller-McNicoll, Sophie Marianne Korn, Andreas Schlundt

## Abstract

AT-rich interacting domain (ARID)-containing proteins, Arids, are a heterogeneous DNA-binding protein family involved in transcription regulation and chromatin processing. For the member Arid5a, no exact DNA-binding preference has been experimentally defined so far. Additionally, the protein binds to mRNA motifs for transcript stabilization, supposedly through the DNA-binding ARID domain. To date, however, no unbiased RNA motif definition and clear dissection of nucleic acid-binding through the ARID domain have been undertaken. Using NMR-centered biochemistry, we here define the Arid5a DNA preference. Further, high-throughput in vitro binding (RBNS) reveals a consensus RNA-binding motif engaged by the core ARID domain. Finally, transcriptome-wide binding (iCLIP2) reveals that Arid5a has a weak preference for (A)U-rich regions in pre-mRNA transcripts of factors related to RNA processing. We find that the intrinsically disordered regions (IDR) flanking the ARID domain modulate the specificity and affinity of DNA-binding, while they appear crucial for RNA interactions. Ultimately, our data suggest that Arid5a uses its extended ARID domain for bi-functional gene regulation and that the involvement of IDR extensions is a more general feature of Arids in interacting with different nucleic acids at the chromatin-mRNA interface.

## Introduction

Among the large number of DNA-binding proteins (DBPs), ARIDs compose a distinct family of nuclear proteins with manifold functions in cellular processes alongside transcriptional regulation (reviewed in ^1,2^). ARID proteins are classified with respect to their shared DNA-binding domain, named AT-rich interactive domain (ARID), reflecting the supposed preference for AT-rich DNA^3,4^. Beyond that, ARID-containing proteins – further referred to as ‘Arids’ for the sake of clear distinction from the ARID domain – are diverse in size and domain architecture, based on which the 15 known human Arids are divided into seven sub-families^5^. All ARID domains share a conserved fold, comprising a minimal core structure of six α-helices (H1 to H6, **Fig. 1**), with H3/4 and H5 forming a central helix-turn-helix (HTH) motif, a widespread DNA-binding unit of DNA-binding domains (DBDs)^5-7^. Turn-containing motifs, similar to the HTH, are in principle also capable of recognizing double-stranded (ds)RNA^8^. It is thus not surprising that the general ability of nucleic acid-binding proteins to interact with both DNA and RNA (DRBPs) is conceived more widespread than previously thought^9^. Still, most DRBPs are assumed to exploit distinct domains to interact with DNA and RNA, respectively, as e.g. known for Sox2^10^ and SAFB proteins^11^. Yet, certain domains, such as the zinc finger motifs, were early found to interact with both types of nucleic acids, e.g. described for the *Xenopus laevis* protein TFIIIA^12^.

**Fig. 1:**
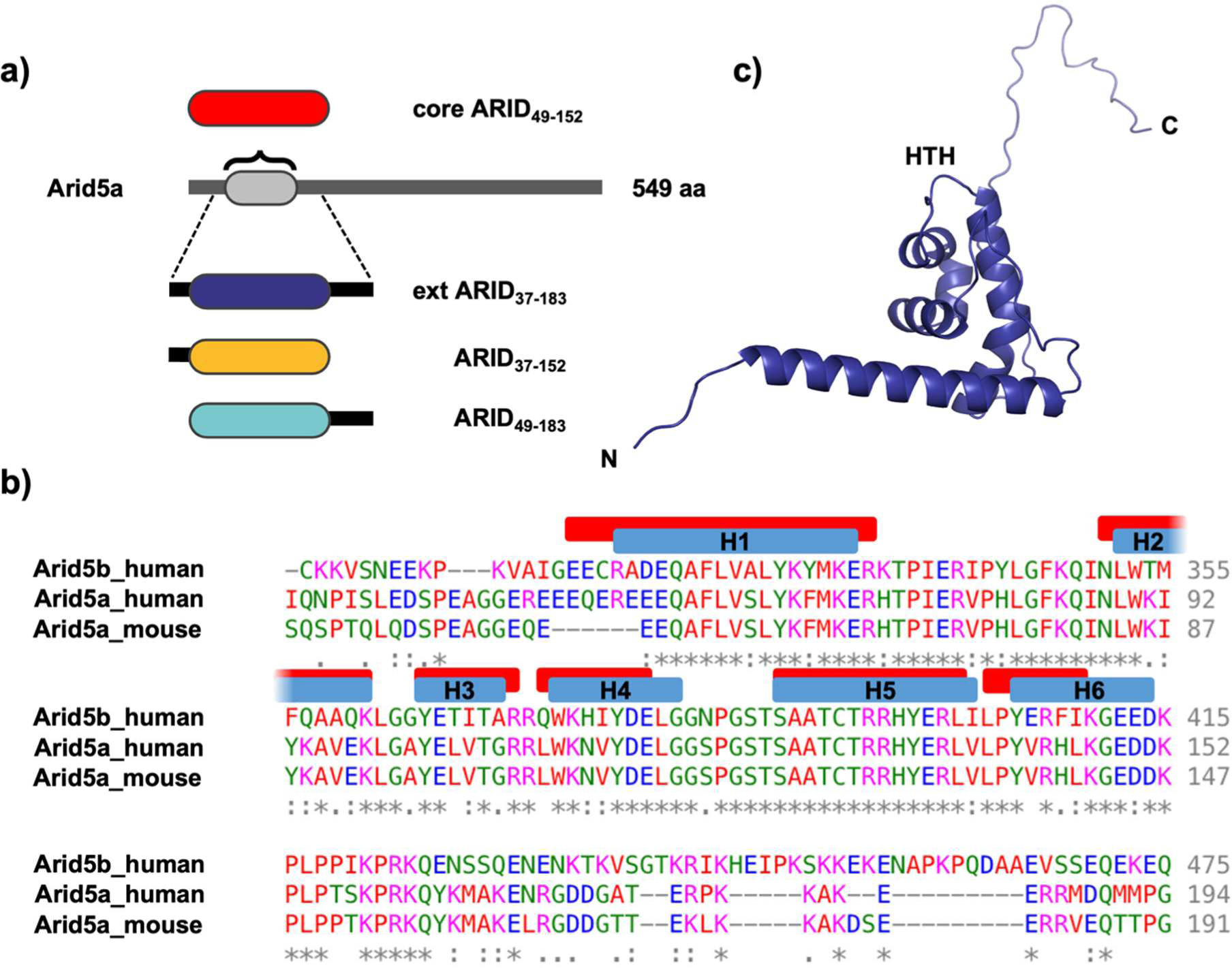
The Arid5a ARID domain is a minimal ARID core motif. **a)** Domain architecture of full-length (fl)-Arid5a and overview of ARID domain boundaries used in this study. **b)** Comparison of human and mouse Arid5a ARID sequences with the human Arid5b ARID, as obtained by Clustal Omega^30^. The ARID secondary structure elements (helices H1 to H6) are indicated above the sequence for human Arid5b (blue, PDB 1IG6) and human Arid5a (red^31^). For full sequence alignment of Arid5a human with mouse see **Suppl. Fig. 2**. **c)** Structural model of the Arid5a core ARID domain as derived from a RoseTTAFold^32^ run with the sequence of ARID_37-183_.

Arid5a is the only Arid representative described as capable of binding both RNA and DNA^13^. The Arid5 family members 5a and 5b share the least conserved domain architecture among Arids. Their ARID domains, however, are 73% identical^5^. The large divergence of Arid5a and 5b reflects distinct functions: Arid5b is categorized as a transcriptional coactivator with essential roles in adipogenesis and liver development, involving chromatin interaction^14^. The existing high-resolution information for an Arid5b ARID-DNA complex has been obtained with the supposedly specific dsDNA consensus motif 5’-AATA[CT]-3’^15,16^. However, the motif has merely been questioned in single-nucleotide exchanges and, more importantly, motif expansions have not been tested. At the same time, a possible capability of the Arid5b ARID domain to interact with (ds)RNA has not been investigated, as is true for all other Arids.

Arid5a, significantly smaller in size, has been classified mainly as a transcriptional repressor^17^, e.g. for nuclear hormone receptors^18^. On the other hand, Arid5a is thought to function actively in the transcription of specific genes and support gene de-repression by histone acetylation together with Sox9^19^. In 2013, Arid5a was termed an RNA-binding protein (RBP), stabilizing the mRNA of *Il-6*^13^, thus counteracting the degradation mediated by the regulatory RBPs Regnase-1 and Roquin^20^. Follow-up work suggested additional targets of Arid5a in an immunological context, among them *Stat3*^21^ and *Ox40*^22^, soon categorizing the protein as pro-inflammatory factor. In these studies, the ARID domain is claimed to interact with particular RNA stem-loop structures, known to exist in *Stat3*^21^, *Ox40*^23,24^ and possibly also the *Il-6* 3’-untranslated region (UTR), suggesting shape-specific recognition of RNA *cis*-regulatory elements similar to Regnase-1 and Roquin^23,25-27^. Though RNA recognition has indirectly been attributed to the Arid5a ARID domain in mice^21^, a direct proof for its interaction with RNA is still missing. At the same time, we have no insight how Arid5a uses its ARID domain to distinguish between specific DNA- and RNA-binding, and whether flanking regions are involved.

Arid5a was initially found differentially expressed in tissues unrelated to the adaptive immune system, but with a clear nuclear localization, in line with transcriptional regulation^17^. Recent work has extended both findings with the protein being able to shuttle upon lipopolysaccharide stimulation in immune cells^28^, a prerequisite for transcript protection against cytoplasmic nucleases. It remains unexplored how RNA motif preferences of the Arid5a ARID domain are related to this, but they should exist independent of cellular localization. There is to date no systematic study identifying the transcriptome targeted by Arid5a independently of its immunological role.

In Arids, the so-called core ARID can appear as N- and/or C-terminally extended domain (**Fig. 1**), i.e. additional helices (H0, H7) or intrinsically disordered regions (IDRs) enlarge the interface with nucleic acids, and likely modify preferences. The strength of IDRs, modifying the function and specifics of DBPs, has recently been brought up for transcription factors (TFs), many with a previously unknown affinity to RNA mediated through their IDRs^29^. While well conceivable as a more general feature, e.g. for compartmentalizing (co)-transcriptional processes, no structural proofs exist for a simultaneous or mutually exclusive interaction of protein domains with RNA and DNA. Similarly, the lack of high-resolution ARID structures with DNAs – with only few exceptions – has hindered us from identifying concepts of specific target recognition through core domains and in combination with flanking regions. As such, most motifs assigned to individual Arids are derived from genetic studies or do not unambiguously define the ARID domain as responsible for interactions. And, the currently known studies have not addressed binding of RNAs by Arids.

We here present a systematic analysis of the Arid5a ARID domain towards specific DNA- and RNA-binding. Using a combination of nuclear magnetic resonance (NMR) and electrophoretic mobility shift assays (EMSAs), we compare nucleic acid recognition of the core with the IDR-extended ARID. Our work provides unambiguous proof for the dual nucleic acid recognition by the domain. We provide in-depth evidence for its preference towards specific AT-DNA motifs, while RNA Bind-n-Seq (RBNS) reveals a preference for an unexpected CAGGCAG consensus motif, accompanied by a general preference for AU-rich motifs (**Data Table 1**). We find that the ARID-flanking IDRs strongly modulate affinity for complex RNA and non-specific DNA sequences. We show that Arid5a exists in the nucleus under unstressed conditions and perform the first individual-nucleotide resolution UV crosslinking and immunoprecipitation

(iCLIP2) experiment to map Arid5a binding sites throughout the transcriptome and identify an *in vivo* (A)U-rich consensus target RNA motif. We find Arid5a to bind RNA-processing related nascent transcripts. While we suggest Arid5a to mainly exert DBP functions, we show that extended ARID domains of other Arids have a similar capacity to interact with RNA. Thus, we stress the idea of Arids as more general dual nucleic acid-binding proteins. We suggest an essential role of the ARID-extending IDRs in nucleic acid recognition, in particular for – but not restricted to – Arid5a.

## Material and methods

### *Arid* protein construct design and mutagenesis

Human Arid5a ARID constructs used in this study were designed and cloned as described previously^31^. In brief, we used different domain boundaries, comprising the minimal ARID core (ARID_49-152_) alone or extended either N- (ARID_37-152_) or C- (ARID_49-183_) terminally, or both (ARID_37-183_), with the numbers representing the natural sequence in the full-length context (**Fig. 1a**). ARID-coding DNA sequences for human Arid1a (residues 999-1132), Arid5b (residues 300-434) and Jarid1a (residues 66-198) were designed to comprise the minimal core ARID plus 18 N- and 20 C-terminal amino acids. They were obtained from Eurofins Genomics, optimized for *E. coli* codon usage, and sub-cloned into the pET24d-derived vector pET-Trx1a (Gunter Stier, EMBL/BZH Heidelberg)^33,34^ by *NcoI/XhoI* restriction and subsequent ligation. Minimal core ARID domains were generated using the respective oligonucleotides listed in **Suppl. Tab. 1**.

Arid5a ARID point mutations, in either the minimal ARID_49-152_ or extended ARID_37-183_ context, were introduced by site-directed mutagenesis (**Suppl. Tab. 1 and 2**). Constructs with multiple non-adjacent mutations were cloned in subsequent steps. A gene encoding for murine full-length Arid5a (fl-Arid5a) (Eurofins Genomics) was cloned into the vector pEGFP-N1 (CLONTECH^®^) (**Suppl. Tab. 1 and 2**) to obtain Arid5a-GFP, for transfection and imaging in human and murine cell lines. Cloning was performed via Gibson assembly^35^. Briefly, the PCR linearized pEGFP-N1 and fl-Arid5a with homologous 5’- and 3’-ends were mixed in the reaction, incubated for 60 min at 50°C and transformed into *E.coli* Dh5α.

To create an Arid5a production vector for a recombinant ARID domain with Strep-tag for RBNS experiments, we amplified the *Arid5a* gene from pET-Trx1a_ARID_49-152_ and cloned it into pET_TRX_Bsa_StrepTag-N via Golden Gate Assembly^36^ using *Bsa*I restriction sites (**Suppl. Tab. 1**). The resulting fusion protein then contains a His_6_-Tag followed by a thioredoxin-tag (TRX), a TEV cleavage site, a Twin-Strep-tag (IBA Lifesciences, Göttingen) and ARID_49-152_. The amino acid sequences are listed **in Suppl. Tab. 6.**

### Protein expression and purification

Proteins were expressed and purified as described recently^31^, with an additional cation-exchange chromatography (CEX) step. An ENrich^TM^ S 5x50 column (Bio-Rad) was equilibrated with low-salt (LS) buffer (20 mM Bis-Tris, 40 mM NaCl, 2 mM DTT, 0.02% NaN_3_, pH 7.2) and subsequently loaded with size-exclusion (SEC) purified protein, buffer-exchanged to LS-buffer in Amicon^®^ ultra centrifugal filters (MWCO: 3 kDa)and concentrated to 5 ml. Pure protein was eluted with a gradient of 0-50% high-salt (HS) buffer (20 mM Bis-Tris, 2 M NaCl, 2 mM DTT, 0.02% NaN_3_, pH 7.2) and a flow rate of 1.25 ml/min. Pure protein (determined by SDS-PAGE) was pooled and re-buffered (20 mM Bis-Tris, 150 mM NaCl, 2 mM TCEP, 0.02% NaN_3_, pH 6.5) in Amicon^®^ ultra centrifugal filters (MWCO: 3 kDa), and subsequently used for NMR, EMSA and RBNS experiments.

### DNA ligand constructs

All DNA oligonucleotides used in this study were obtained from Sigma-Aldrich. dsDNA was obtained through annealing of complementary strands (5 min at 98°C followed by cooling down to room temperature). An overview of herein used DNAs is given in **Suppl. Tab. 3 and 5**.

### RNA *in vitro* transcription

Unlabelled RNAs from 15 nt in length and longer were produced by in-house optimized *in vitro* transcription (IVT) and purified either from a linearized plasmid or from annealed oligonucleotides (**Suppl. Tab. 4**) as described in^37^. Briefly, plasmid DNA was linearized with *HindIII* prior to IVT by in-house expressed T7-RNA polymerase. Alternatively, complementary oligonucleotides (Sigma-Aldrich) were annealed and used as templates for IVT. RNAs from preparative-scale (10 to 20 ml) transcription reactions (4 h at 37°C) were precipitated with 1.5 volumes 2-propanol overnight at −20°C. RNAs were separated on 12-18% denaturing polyacrylamide gels and visualized by UV shadowing. The excised RNA-fragments of expected length were eluted into 0.3 M NaOAc overnight and subsequently washed, concentrated, and buffer-exchanged to the experimental buffer.

RNAs below 15 nt in length (**Suppl. Tab. 5**) were obtained from Dharmacon Horizon in 1-µmol-scale quantities, deprotected and desalted. Each RNA was dissolved in the respective volume of ddH_2_O to a final concentration of 3 mM.

### *In vitro* transcription of the RBNS input pool

As template a T7 promoter-containing oligonucleotide was annealed to an equimolar quantity of RBNS T7 template oligonucleotide (a random 20-mer flanked by partial Illumina primers). 500 fmol template were transcribed overnight at 37°C with 200 mM Tris-HCl pH 8.0, 20 mM magnesium acetate, 8% (v/v) DMSO, 20 mM dithiothreitol (DTT), 20 mM spermidine, 4 mM nucleoside triphosphates (NTP) (each) and self-made T7 RNA-Polymerase. The RBNS pool was purified by PAGE (polyacrylamide gel electrophoresis). Oligonucleotide sequences are given in **Suppl. Tab. 7**.

### NMR spectroscopy

NMR experiments were performed at the Frankfurt BMRZ using Bruker Avance III/Avance Neo spectrometers of 600 and 900 MHz proton Larmor frequency, equipped with cryogenic probes and using Z-axis pulsed field gradients. All measurements containing protein were performed at 298 K. Topspin versions 3 and 4 were used for data acquisition and processing. Graphical plots of spectra were created using the program NMRFAM-Sparky^38^ version 1.470.

Titrations were performed by preparing two initial samples: i) a protein apo sample and ii) a sample comprising protein in the presence of the maximum DNA/RNA concentration. All intermediate titration points were mixed from those samples subsequently (from high to low) to avoid side effects of protein dilution. For each sample we monitored protein peaks by recording ^15^N-(TROSY)-HSQCs and DNA/RNA imino peaks by acquisition of 1D imino proton spectra. For HSQC-spectra, we typically recorded 128 and 2048 points in the indirect ^15^N and ^1^H direct dimensions, respectively, with spectral widths of 32 ppm (offset at 116.5 ppm) and 16 ppm. For 70 µM samples used in titrations, we recorded 32 scans per increment, while 40 scans per increment were recorded for 50 µM samples. For the DNA 1D imino proton spectra, we recorded a second set of experiments, where the DNA concentrations were kept constant at 80 µM and the protein concentration varied (10 µM, 20 µM, 40 µM, 80 µM). Spectra were recorded with 8192 points and 512 scans for 13merAT and 2560 points and 256 scans for 13merGC. The spectral width was set to 23.5 ppm and 21 ppm for 13merAT and 13merGC, respectively. Analysis of spectra and quantification/plotting of CSPs from titrations were performed in the CCPNMR Analysis 2.5 software^39^. Significance of CSPs was defined as above average plus one standard deviation (SD), if not indicated differently.

For the assignment of imino protons in the 13merAT, we recorded a ^1^H-^1^H-NOESY at 278 K with a spectral width of 22 and 15 ppm and 4096 and 266 points for the direct and indirect proton dimensions, respectively. The mixing time was set to 300 ms. Based on this we transferred the assignment to 298 K in a peak-traceable temperature series (**Suppl. Fig. 5**).

### Structures and structure models

Structural models of Arid5a were generated with RoseTTAfold^32^ using residues 37-183 from the sequence deposited in Uniprot^40^ under ID Q03989. The 13merAT dsDNA was modelled in the program Avogadro^41^ from its primary sequence as B-DNA. PyMOL (The PyMOL Molecular Graphics System, Version 2.0 Schrödinger, LLC.) was used to align both the Arid5a and the 13merAT dsDNA model individually to the structure of Arid5b ARID in complex with DNA (PDB 2OEH^16^). The aligned models were than manually arranged to each other by means of a slight positional adjustment to create a model of the extended Arid5a ARID domain binding to DNA.

### EMSA

To decipher the interaction of RNA/DNA and protein, we used electrophoretic mobility shift assays (EMSA) with radioactively labelled RNA (rEMSA) and fluorescently labelled DNA. The RNA was *in vitro* transcribed with T7-RNA polymerase and labelled with γ-^32^P according to a protocol by Nahvi and Green^42^. We used 30 pmol of RNA, which was dephosphorylated at the 5’-end with 3 µl of Quick-CIP (5000 U/µl, NEB) in a total volume of 20 µl according to manufactureŕs instructions. Next we performed a phenol/chloroform extraction and precipitated the RNA with ethanol and sodium acetate in the presence of 20 µg glycogen for 30 min at -20°C. The precipitated RNA was pelleted for 15 min at 16,000 g at 4°C. The pellet was resuspendend in 10 µl ddH_2_O from which 5 µl were used for the ^32^P-labeling. Therefore 1.5 µl γ-^32^P-ATP (10 pmol, Hartmann Analytic), 2 µl T4-PNK buffer (NEB), 2 µl T4-PNK (10 U/µl, NEB) and 9.5 µl H_2_O_MQ_ were added. The reaction was incubated for 60 min at 37°C to allow phosphorylation followed by 10 min at 80 °C to inactivate the kinase. To finally purify the radioactively-labelled RNA we used *NucAway Spin Columns* (Thermo Fisher Scientific) according to manufacturer’s instructions. Finally, the RNA was refolded (4 min 95°C, cooled down on ice water) and diluted to a final volume of 400 µl and stored at -20°C.

For the fluorescently labelled DNA, complementary DNA oligonucleotides were used (**Suppl. Tab. 3**), with one oligonucleotide 5’-labelled with fluorescein- (FAM) and the other one unlabelled. Complementary oligonucleotides were mixed in a 1:1 ratio and heated to 9°C for 5 min before cooling down to allow for the annealing of dsDNA.

EMSA reactions were prepared in a final volume of 20 µl. Therefore, we mixed 0.6 µg of yeast tRNA (Roche), 10 mM MgCl_2_, Arid5a-buffer (20 mM Bis-Tris, 150 mM NaCl, 2 mM TCEP, 0.02% NaN_3_, pH 6.5) and respective amounts of protein. Finally, 2 µl labelled RNA or DNA were added and the reaction incubated for 10 min at room temperature (22-24°C). Immediately before loading 10 µl onto a 6-% polyacrylamide gel, 3 µl of loading buffer were added. Gel-electrophoresis was run for either 40 min at 80 V for DNA or 80 min at 80 V for RNA. The gels were imaged with a Typhoon Imager (GE Healthcare) either in the glass plates (for DNA) with a laser at 488 nm excitation and an emission filter at 520 nm or dried and indirectly imaged by phosphor imaging (for RNA).

### RNA Bind-n-Seq (RBNS)

An RBNS assay was performed with the Twin-Strep-tagged ARID_49-152_ domain and a randomized input RNA pool based on reference^43^. The protein was equilibrated in binding buffer (25 mM Tris-HCl, pH 7.5, 150 mM KCl, 3 mM MgCl_2_, 0.01% Tween, 500 µg/ml BSA, 1 mM DTT) at three different concentrations (0.25, 1, 5 µM) for 30 min at 4°C. Next, the RNA was folded by snap-cooling and added to a final concentration of 1 µM with 40 U Ribonuclease Inhibitor (moloX, Berlin) and incubated for 1 h at room temperature. A pulldown was performed by incubating the RNA/ARID mixture with 1 μl of washed MagStrep “type3” XT beads (IBA Lifesciences, Göttingen) for 1 h at 4°C. Subsequently unbound RNA was removed by washing three times with wash buffer (25 mM Tris-HCl pH 8.0, 150 mM KCl, 60 μg/ml BSA, 0.5 mM EDTA, 0.01% Tween). Afterwards, the RNA/ARID complexes were eluted twice with 25 µl of Elution Buffer (wash buffer containing 50 mM biotin). RNA was extracted with the Zymo RNA Clean & Concentrator-5 kit (Zymo Research, Freiburg) according to the manufacturer’s instructions. The extracted RNA was reverse transcribed into cDNA, amplified by PCR to add Illumina adapters (**Suppl. Tab. 7**) and an index for each concentration (**Suppl. Tab. 8**), and subjected to deep sequencing (GENEWIZ, Leipzig).

Next-generation sequencing data were analyzed using the RBNS pipeline as described in^44^, available at https://bitbucket.org/pfreese/rbns_pipeline/overview. The sequence context was analyzed using a self-written Python script. This searches for a given motif (in this case the enriched kmers) in each read of the sequence and generates the upstream and downstream sequence logos of the given sequence. Logos are then calculated from this, which are corrected by the composition of the bases (background) in the input pool.

### Culturing and transfection of P19 cells

Murine P19 WT cells were cultivated on 10-cm culture dishes pre-coated with 0.1% gelatin (in PBS) under humidified condition at 5% CO_2_ and 37°C in DMEM GlutaMAX Medium, supplemented with 10% (v/v) heat inactivated fetal bovine serum and 100 µg/ml penicillin-streptomycin (all Gibco™, Thermo Fisher Scientific). P19 WT cells were transfected with 4 µg plasmid DNA in 10-cm plates using the jetOPTIMUS^®^ Transfection reagent (Polyplus) according to the manufacturer’s instructions. The cells were harvested after 24 h of incubation.

### Confocal microscopy

For GFP- and immunofluorescence (IF) microscopy, P19 cells were grown on pre-coated 10 mm glass coverslips in 10-cm plates. The coverslips were transferred into a 24-well plate and washed with 1x PBS. After removing the PBS, cells were fixed with 4% paraformaldehyde (in PBS; Thermo Fisher Scientific) for 20 min at room temperature. Fixed cells were washed twice with 1x PBS and then permeabilized in permeabilization buffer (5% BSA, 0.1% Triton in 1x PBS) for 30 min. Mouse anti-G3BP1 antibody (Abcam, ab56574) was diluted in blocking buffer (5% BSA in 1x PBS) at 2 µg/ml final concentration and incubated for 16 h overnight at 4°C in the dark as a cytoplasmic marker. The coverslips were washed twice with 1x PBS and incubated with the secondary antibody (donkey anti-mouse coupled to Alexa Fluor 594, Abcam; 1:500 in blocking buffer) for 60 min at room temperature in the dark. After washing the coverslips twice with 1x PBS the DNA was stained with Hoechst 34580 (Thermo Fisher Scientific) at a final concentration of 5 µg/ml in TBST (Tris-buffered-saline with 0.1% Tween-20) for 30 min at room temperature in the dark. After a final wash, the coverslips were dried and mounted on ProLong™ Diamond Antifade Mountant (Thermo Fisher Scientific P36961).

Images were acquired with confocal laser-scanning microscope (LSM780; Zeiss) with a Plan-Apochromat 63x 1.4 NA oil differential interference contrast objective M27 using the Zen 2012 (black edition; 8.0.5.273; ZEISS). Fluorescence signal was detected with an Argon laser (GFP – 488 nm, G3BP1/Qasar – 594 nm and Hoechst – 405 nm). Fiji was used to crop the pictures with the Image crop function and to add the scale bars^45^.

### iCLIP2 of full-length Arid5a-GFP

iCLIP experiments were performed using the iCLIP2 protocol^46^ with minor modifications. For each replicate, two 15-cm dishes of P19 cells grown to 60% confluence were transfected with 15 µg of Arid5a-GFP plasmid DNA. After 24 h, the cells were washed with ice-cold PBS, irradiated with 250 mJ/cm^2^ UV light at 254 nm (CL-1000, UVP) and harvested by scraping and centrifugation. Following lysis and partial digestion with RNase I (Thermo Fisher Scientific, AM2294), immunoprecipitation of Arid5a-GFP was performed using a goat anti-GFP antibody (MPI-CBG, Dresden, Germany) coupled to Dynabeads™ Protein G (Thermo Fisher Scientific, 10002D). Co-purified, crosslinked RNA fragments were dephosphorylated at their 3’-ends using T4-PNK (NEB, M0201S) and ligated to a pre-adenylated 3’-adapter (L3-App, **Suppl. Tab. 9**). To visualize protein-RNA complexes, RNA fragments crosslinked to Arid5a-GFP were labelled at their 5’ ends using T4-PNK and γ-^32^P-ATP (Hartmann Analytic). Samples were run on a Nu-PAGE 4-12% Bis-Tris Protein Gel (Thermo Fisher Scientific, NP0335BOX), transferred to a 0.45 µm nitrocellulose membrane (GE Healthcare Life Science, 10600002) and visualized using a Phosphorimager. Regions of interest were cut from the nitrocellulose membrane (95 kDa to 180 kDa), and RNA was released from the membrane using Proteinase K (Roche, 03115828001). RNA was purified using neutral phenol/chloroform/isoamylalcohol (Ambion, AM9722) followed by chloroform (Serva, 39554.02) extraction, and reverse transcribed using SuperScript III™ (Life Technologies, 18080-044) and a short RT primer (**Suppl. Tab. 9**). cDNA was cleaned up using MyONE Silane beads (Life Technologies, 37002D) followed by ligation of a second adapter containing a bipartite (5-nt + 4-nt) unique molecular identifier (UMI) as well as a 6-nt experimental barcode^46^ (Lclip2.0 adapter, **Suppl. Tab. 9**). iCLIP2 libraries were pre-amplified with 6 PCR cycles using short Solexa primers P5 and P3 (**Suppl. Tab. 9**) and then size-selected using the ProNex Size-Selective Purification System (Promega, NG2001) in a 1:2.95 (v/v) sample:bead ratio to eliminate products originating from short cDNAs or primer dimers. The size-selected library was amplified for 6 cycles using long Solexa primers P5 and P3 (**Suppl. Tab. 9**), and primers were removed using the ProNex Size-Selective Purification System (Promega, NG2001) in a 1:2.4 (v/v) sample:bead ratio. Purified iCLIP2 libraries were sequenced on a NextSeq 500 System (Illumina) using a NextSeq^®^ 500/550 High Output Kit v2 as 92-nt single-end reads, yielding between 18 and 21 million reads.

### iCLIP2 analysis

iCLIP2 data were processed as described in^47^. In brief, quality control was done using FastQC (version 0.11.9) (https://www.bioinformatics.Babraham.ac.uk/projects/fastqc/). Read were de-multiplexed according to the sample barcode on positions 6 to 11 of the reads using Flexbar (version 3.5.0, ^48^) using non-default parameters: flexbar --adapter-seq AGATCGGAAGAGCGGTTCAG --adapter-min-overlap 1 --min-read-length 15 --length-dist -- umi-tags. Flexbar was also used to trim UMI and barcode regions as well as adapter sequences from read ends requiring a minimal overlap of 1 nt of read and adapter. UMIs were added to the read names and reads shorter than 15 nt were removed from further analysis. The downstream analysis was done as described in ^46^. Reads were mapped with STAR (v2.7.3a)^49^ with non-default parameters: STAR --alignEndsType Extend5pOfRead1 -- outFilterMismatchNoverReadLmax 0.04 --outFilterMultimapNmax 1. Genome assembly (GRCm38.p6) and annotation of GENCODE (release M25)^50^ were used.

Reads directly mapped to the chromosome ends were removed, as they do not have an upstream position and, no crosslink position can be extracted using Samtools (v1.10)^51^, bedtools (v2.29.2)^52^. PCR duplicates were removed using UMI-tools (v1.1.2) with non-default parameters: umi_tools dedup --method unique –random-seed=100. To extract the crosslink sites the bam files were first converted into bed files shifting one base upstream using bedtools (v2.29.2) bamtobed and shift. Then only the first nucleotide was kept and the positions separated by the strand information using bedtools (v2.29.2) genomecov. Processed reads from three replicates were merged prior to peak calling with PureCLIP (version 1.3.1) ^53^ using a minimum transition probability of 1%. Significant crosslink sites (1 nt) were filtered by their PureCLIP score, removing the lowest 1% of crosslink sites. The remaining sites were merged into 7-nt wide binding sites using the R/Bioconductor package BindingSiteFinder (version 2.0.0), filtering for sites with at least 2 positions covered by crosslink events. Only reproducible binding sites were considered for further analyses, which had to be supported by two out of three replicates. Binding sites were overlapped with gene and transcript annotations obtained from GENCODE (release 29). Binding sites in intergenic regions were removed from further analysis. Binding sites within protein-coding genes were assigned to the transcript regions, i.e., intron, coding sequence, 3’-UTR or 5’-UTR.

Motif analysis was performed by counting all possible 3-mers in a window of ±50 nt around the center points of all Arid5a binding sites using the R/Bioconductor package Biostrings (version 2.70.1). Heatmap visualization was done separately for 3-mers starting and/or ending on U (**Fig. 7c**) and all other 3-mers (**Fig. 7d**), including *k*-means clustering with *k*=3.

## Results

### Highly conserved ARID domains show distinct DNA-binding preferences

Doubts have evolved over recent years to whether all name giving AT-rich interactive domains of the 15 human Arids share exclusive preference for AT-rich sequences (recently reviewed by Korn and Schlundt^5^). Indeed, controversial sequence preferences reported for a number of Arids gave reasons to unbiasedly probe for individual target sequences^1,54-56^. We thus picked representative ARID domains from three sub-families that had been described to target DNA with different sequence preferences (**Fig. 2a**). While some literature describes Arid1a to bind DNA non-specifically through its ARID domain, the ARID domains of Arid5b and JARID1a are suggested to be specific for AT- and GC-rich dsDNA, respectively^1,16,56^. We used fluorescently labelled AT- or GC-rich dsDNA to monitor preferences of these ARID domains in EMSAs (**Fig. 2b and Suppl. Fig. 1**). Interestingly, the ARID domains of Arid1a and JARID1a are less specific for AT-rich DNA than the ARID domain of Arid5b (see also **Suppl. Fig. 1**). Furthermore, and in line with multiple studies^54,55,57^, the Arid1a ARID domain displays similar affinity for a GC-rich dsDNA, supporting its non-specificity for DNA. In summary, the data argue against ARID domains as exclusive AT-binders and raise the need to carefully *de novo* define and interpret available consensus motifs for the individual domains despite their highly conserved fold.

**Fig. 2:**
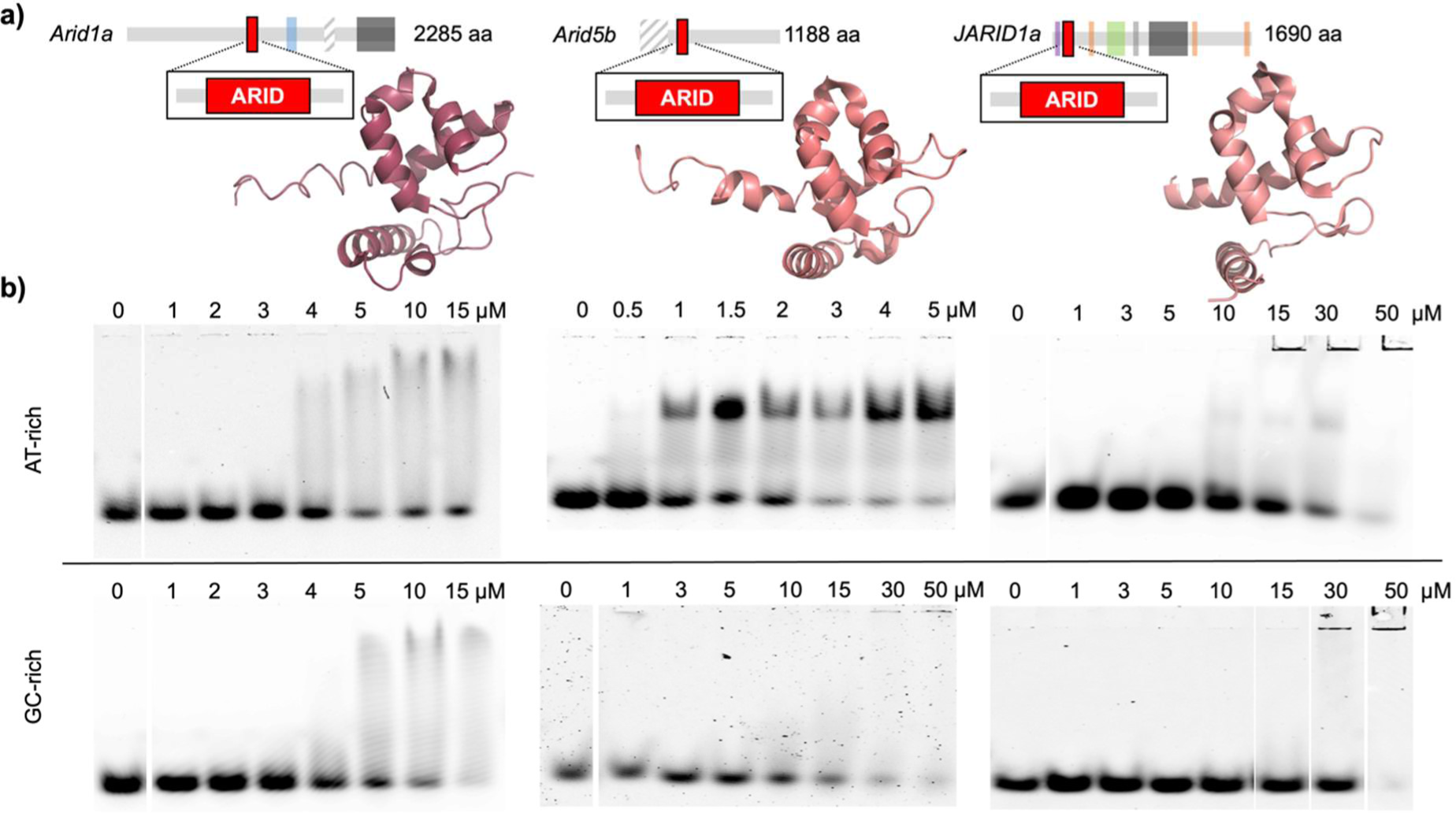
DNA-binding preferences of ARID domains from three different human Arid sub-families. **a)** Domain architecture of Arid1a, Arid5b and Jarid1a, with the ARID domain indicated by a red box and their structures depicted below (Arid1a: PDB 1RYU^58^; Arid5b: PDB 1IG6, unpublished; and Jarid1a: PDB 2JXJ^59^). Other annotated domains are HIC1 (light blue), BAF250-C (dark grey), JMJN (purple), JMNC (green), PHD (orange) and ZnF (light grey). Merely predicted domains are striped black and white. **b)** The DNA-binding preference of extended ARID domains comprising the minimal core ARID plus 18 N- and 20 C-terminal residues were studied by EMSAs with either 13merAT or 13merGC fluorescently labelled dsDNA (**Suppl. Tab. 3**).

### The Arid5a ARID domain uses an extended binding interface with AT-rich DNA

Because of the above-described variance in the DNA-binding preferences of ARID domains, we first decided to investigate the Arid5a ARID domain’s sequence preference. Although 9mer dsDNA sequences were sufficient for binding, we observed a minor increase in affinity with longer dsDNAs plateauing at 13 bp length (**Suppl. Fig. 3**), and thus used 13mers for our study. In EMSAs, we tested ARID_37-183_ against fluorescently labelled dsDNAs, either GC-rich based on Jarid1a binding to a “CCGCCC” motif^56^ or with variations of a central AT-stretch based on a published motif for the closely related Arid5b ARID^16^ (**Suppl. Tab. 3 and Suppl. Fig. 4**). We find that the Arid5a ARID domain clearly favors AT-rich dsDNA, and a central “AATA” motif is important as evident from apparent affinities around 1-2 µM contrasting the at least 10-fold higher K*_D_* for DNAs without the motif (**Suppl. Fig. 4**).

To investigate differential complex formation on the residue-resolved level, we next used NMR and performed ^1^H-^15^N-heteronuclear single quantum coherence (HSQC) titrations of either 13merAT or 13merGC dsDNA to the extended ARID_37-183_ and plotted the combined chemical shift perturbations (CSPs) over the protein sequence (**Fig. 3a and b**). With this experiment we sought to i) identify the precise interface(s) of the ARID domain with DNA beyond its core fold and ii) spot potential differences in CSP patterns caused by the two dsDNA ligands. The titrations clearly show that ARID_37-183_ binds to both the 13merAT and the 13merGC dsDNA. However, different exchange regimes – fast exchange for 13merGC and intermediate exchange for 13merAT (insets **Fig. 3a**) – support the significantly higher affinity of Arid5a ARID to AT-rich as compared to GC-rich DNA observed in EMSAs (**Suppl. Fig. 4**). Of note, maximum CSPs within core ARID residues are much smaller for 13merGC than for the AT-rich DNA (**Fig. 3a and b**). Interestingly, CSP differences of flanking IDR residues, especially the C-terminal extension (residues 150 – 160), are less pronounced between GC- and AT-rich DNA targets than within the core domain, suggesting the contribution of IDRs to DNA-binding is less or non-specific.

**Fig. 3:**
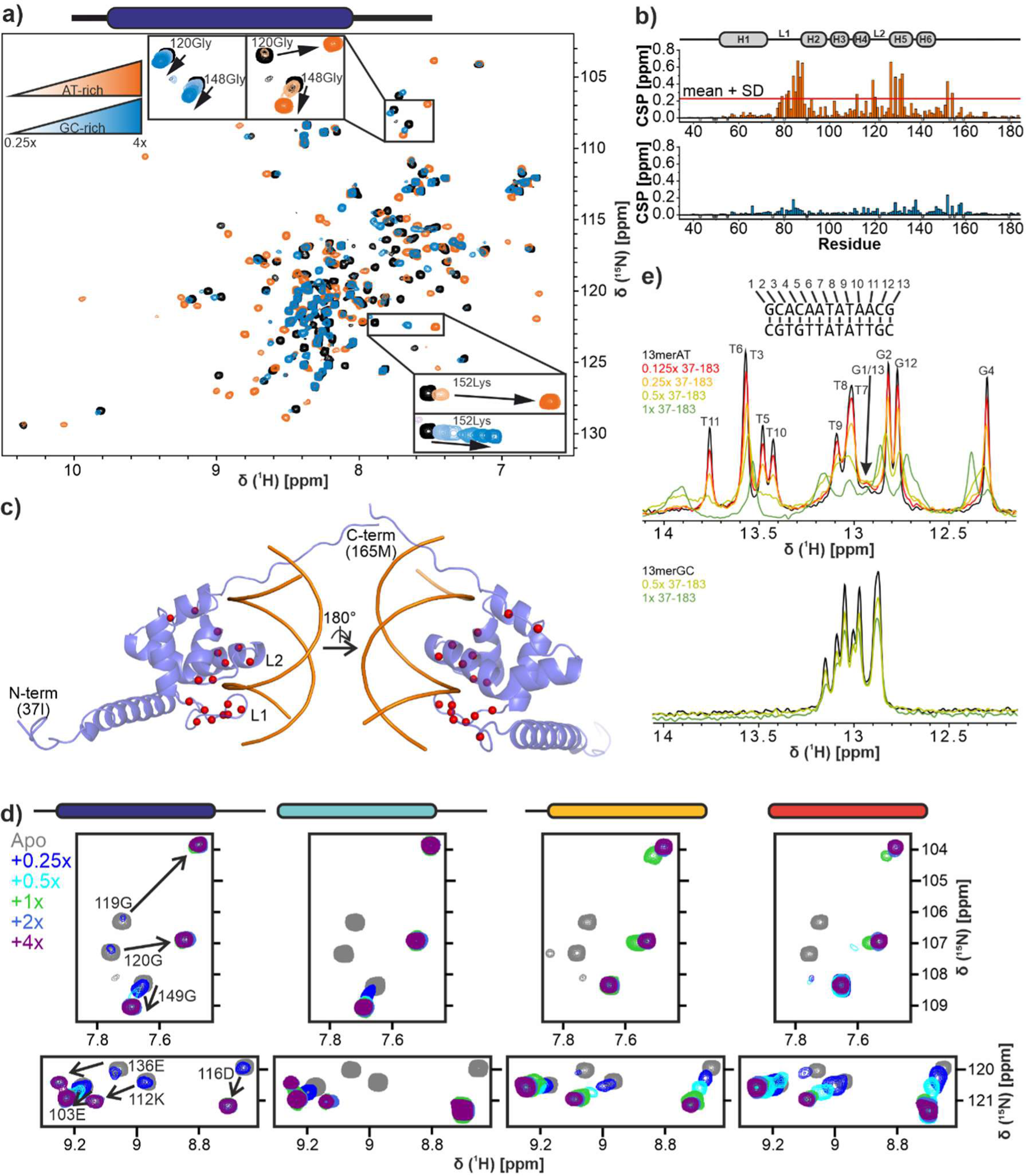
Arid5a interacts with AT-rich DNA through loops in its core ARID and the C-terminal IDR. **a)** ^1^H-^15^N-HSQC overlay of ARID_37-183_ alone and after titration with 4-fold 13merAT (orange) or 13merGC (blue). Insets show all titration points and assignments. **b)** Chemical shift perturbation (CSP) plot of ARID_37-183_ upon titration of 4-fold 13merAT (upper panel) or 13merGC (lower panel). Negative bars in light grey and gray show prolines and unassigned residues, respectively. Significantly shifting peaks, i.e. above mean+SD, are shown in **Suppl. Fig. 7b**. **c)** RoseTTAFold^32^ model of ARID_37-183_ with highest CSPs from **b)** (above mean +1 standard deviation) highlighted in red. For simplification, only residues 37-165 are shown. **d)** ^1^H-^15^N-HSQC zoom-ins of four Arid5a ARID constructs with/without N- and/or C-terminal extension – as indicated above – showing spectra of proteins alone and when titrated with 13merAT dsDNA. **e)** 1D imino proton spectra of 13merAT (upper panel) and 13merGC (lower panel) upon titration of DNAs with ARID_37-183_. See also **Suppl. Fig. 5**.

From the CSP plots we concluded that the Arid5a ARID domain interacts with DNA through residues in loop L1 and the HTH motif (H4-L2-H5) (**Fig. 3b**). This is in good agreement with the reported DNA-binding interface found for other ARID domains^16,58,60,61^ and an R-to-A mutant in murine Arid5a (corresponding to R133 in the human version, see **Fig. 1b**) incapable of DNA-binding^21^. Importantly, our data reveal an additional contribution of residues K152 and L154 within the C-terminal extension. Mapping significant CSPs obtained for the AT-DNA interaction on an Arid5a ARID RoseTTAFold model clearly shows them to cluster in the canonical DNA-binding interface (**Fig. 3c**).

To investigate the potential contribution of both N- and C-terminal extensions to DNA-binding in more detail we created constructs of the ARID domain with either the separate N- (ARID_37-152_) or C-terminal (ARID_49-183_) extension and compared their DNA interaction with 13merAT to the core domain (ARID_49-152_) and the extended ARID_37-183_ (**Fig. 3d and Suppl. Fig. 6 and 7**). In contrast to the N-terminal IDR, the C-terminal extension shifted the ARID-DNA interaction towards an NMR-observed intermediate-to-slow exchange regime (**Fig. 3d and Suppl. Fig. 6**), supported by observable changes in the EMSA patterns (**Suppl. Fig. 8**). The latter does not only support the higher affinities for C-terminally extended ARID constructs (ARID_49-183_ and ARID_37-183_), but also reveals formation of distinct complex bands, indicating compact DNA-protein complexes that are less pronounced in ARID domains devoid of the C-terminal extensions.

To confirm sequence-specific DNA recognition in the 13merAT DNA compared to 13merGC, we titrated increasing concentrations of ARID_37-183_ to the respective DNAs and monitored effects on imino protons (**Fig. 3e**). We undertook a complete assignment of 13merAT imino resonances, which allowed a base pair-resolved analysis (**Suppl. Fig. 5**). In line with the EMSA-observed stable complex formation, we found severe line-broadening for the central AATA-motif in 13merAT, accompanied by CSPs of the neighboring base pairs. In contrast, the 13merGC merely displayed minor line broadening upon ARID_37-183_ addition, evenly distributed over all imino signals. This supports a weak, but non-specific interaction with the GC-rich DNA, driven by electrostatic interactions with the DNA backbone rather than base-specific contacts.

### Mutational studies of Arid5a confirm key residues for DNA-binding

To confirm the ARID DNA-binding interface, we designed protein mutants by replacing selected residues, located either in L1 or the HTH motif, by alanine. Residues were chosen based either on their high CSPs observed in the ARID_37-183_ titration with 13merAT (K85 and Q86) or on literature and sequence comparison to other Arids – especially Arid5b – and their key DNA-binding residues (R78A, R109A, T125A/S126A)^1,5,16^. Mutations were introduced both in the core ARID_49-152_ and extended ARID_37-183_ background, to further elucidate the role of IDRs in this context. ^1^H-^15^N-HSQC spectra of proteins alone and in presence of 2-fold molar excess 13merAT dsDNA were recorded to quantify the effect of single, double and triple mutations on DNA-binding (**Fig. 4 and Suppl. Fig. 9 and 10**).

**Fig. 4:**
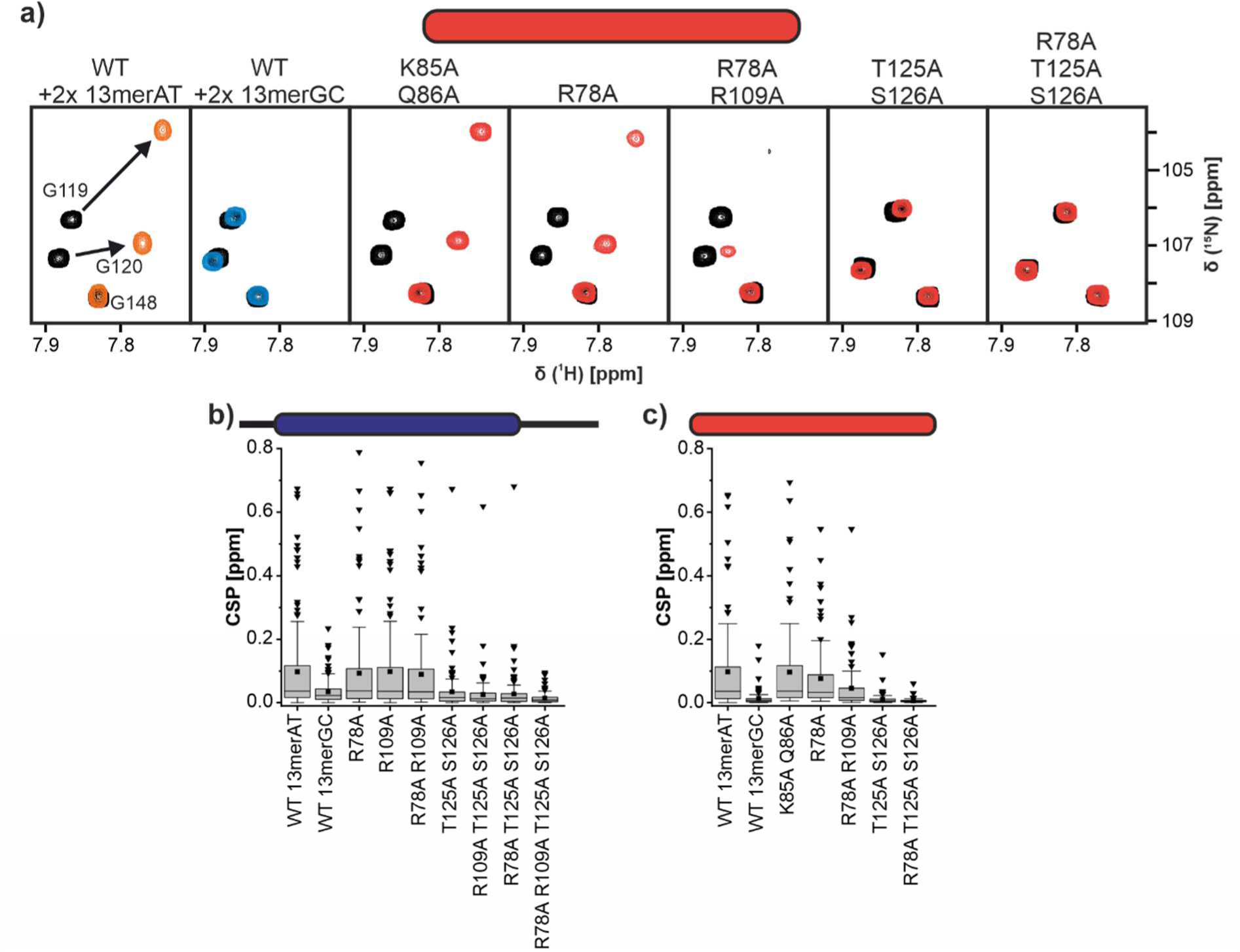
Mutational studies of the Arid5a ARID domain reveal the central core binding residues. **a)** ^1^H-^15^N-HSQC zoom-ins of ARID_49-152_ wild type (WT) and mutants overlaying apo protein spectra (black) with those of samples containing 2-fold 13merAT dsDNA (orange/red) or 2-fold 13merGC dsDNA (blue). **b)** Boxplot of CSP quantifications of ARID_37-183_ WT and mutants upon addition of 2-fold 13merAT dsDNA. **c)** Boxplot of CSP quantifications of ARID_49-152_ WT and mutants upon addition of 2-fold 13merAT dsDNA. For comparison and as a reference each boxplot also shows the CSPs of the WT with 2-fold 13merGC dsDNA. The box represents the interquartile range from the 25^th^ to the 75^th^ percentile with a whisker coefficient of 1.5 for outliers and further outliers shown as black triangles. The median is shown as a horizontal line within boxes and mean values are indicated by black squares. Source data are provided as a Source Data file.

Mutation of T125 and S126, located in the HTH at the transition of loop 2 to helix 5, strongly impaired DNA-binding of the core ARID domain (**Fig. 4a**). This is in line with their expected role in making specific contacts with an AT base pair in the DNA major groove, as suggested by the complex structure of the closely related Arid5b ARID domain with AT-rich dsDNA^15^. Loop 1 mutations (K85A/Q86A) on the other hand – despite high CSPs (**Fig. 3**) – did not inhibit DNA-binding and likely exhibit no crucial DNA contacts. Of note, the effect of DNA-binding mutations was less pronounced in presence of the extending IDRs, evident when comparing global CSPs between ARID_37-183_ and ARID_49-152_ (**Fig. 4b and c**). We thus conclude that the C-terminal extension to the ARID domain can compensate for mutations within the core ARID domain by a general, but non-specific mode of increasing affinity for dsDNA.

### *In vitro* RNA-binding of the Arid5a ARID domain

Arid5a was recently identified to stabilize the *Ox40* mRNA in murine CD4+ T cells by direct interaction with a stem-looped structure in its 3’-UTR, known as an alternative decay element (ADE)^22^. In doing so, Arid5a interferes with controlled degradation of the *Ox40* transcript by the nuclease Regnase through targeting the same *cis*-regulatory element. We wondered if the ARID domain in Arid5a was responsible for the underlying complex formation with the RNA-stem loop and used NMR spectroscopy to observe atom-resolved binding of the ARID domain to the ADE element (**Fig. 5 and Suppl. Fig. 11**). Interestingly, the minimum core ARID_49-152_ showed only marginal interactions, even with a high stoichiometric excess of the 19-nt ADE, (**Fig. 5d and e and Suppl. Fig. 11**). However, similar to DNA-binding, the basic C-terminal extension contributed to an increased ADE interaction (**Suppl. Fig. 11**), suggesting the extension to carry an essential role in Arid5a*-*based mRNA regulation *in vivo*.

**Fig. 5:**
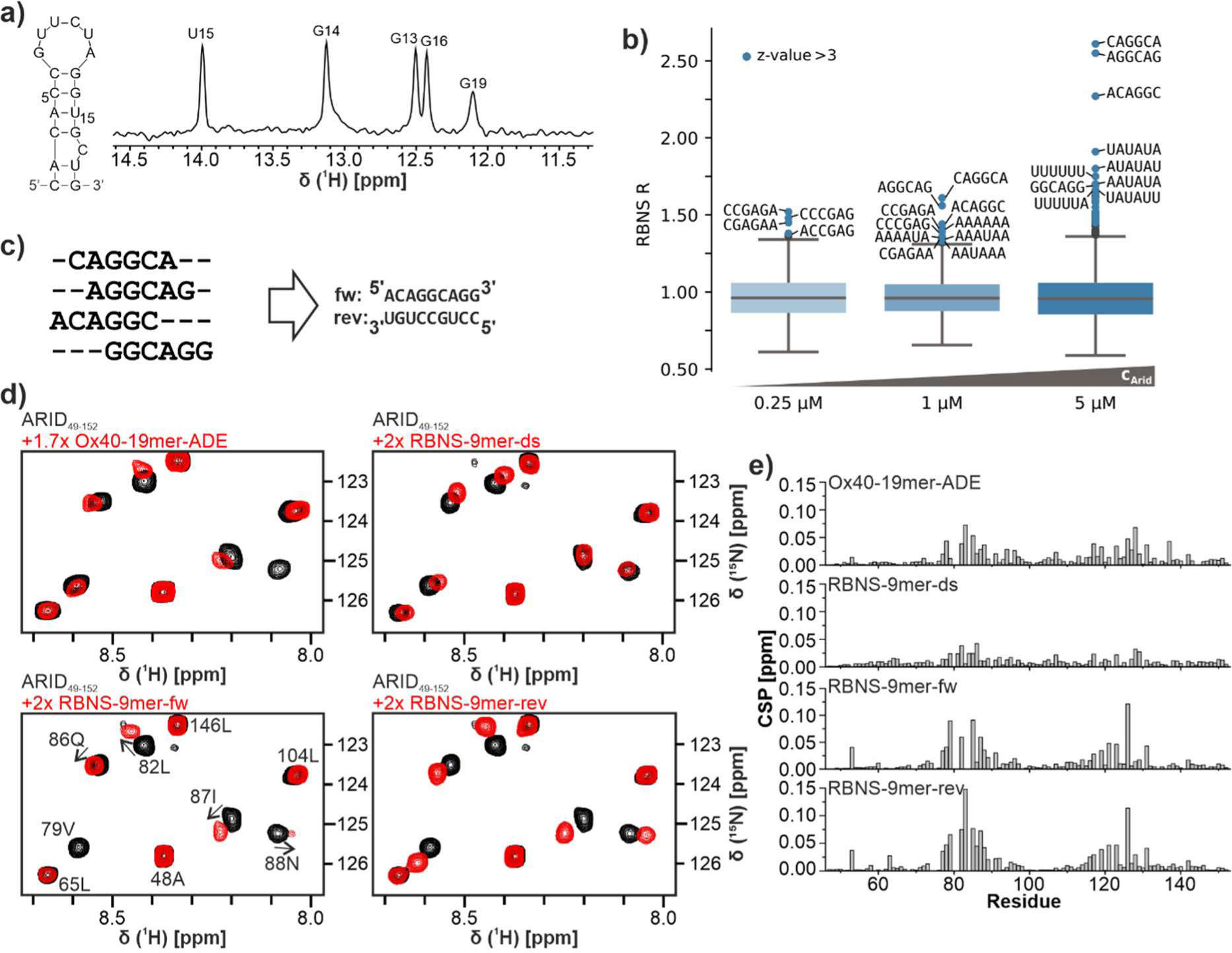
The Arid5a ARID domain binds RNA motifs with moderate affinity. **a)** The *Ox40*-ADE forms a stem-loop element, confirmed by the depicted imino-proton spectrum. Assignments have been transferred from Janowski *et al*., 2016^23^. **b)** Enrichment of all 6-mers at 0.25, 1 and 5 µM ARID_49-152_ concentration. Values greater than three standard deviations above the mean are highlighted blue. For the highest significant ten motif sequences are given **c)** RBNS-based 9mer sequences that can be clustered from the enriched 6mers containing AGGC. **d)** Zoom-ins of ^1^H-^15^N-HSQC spectra of apo ARID_49-152_ overlaid with 1.7-fold molar excess of ADE RNA or 2-fold molar excess of RBNS-9mer RNAs **e)** CSP plots of ARID_49-152_ upon addition of *Ox40*-ADE or RBNS-9mer RNAs as shown in d).

While our data are the first structural proof of a direct ARID-RNA interaction, we were surprised by the observed moderate binding affinity. To this end, we decided to set up an unbiased search for a general consensus RNA target motif of the core ARID fold, which had not been undertaken prior to this study. We thus performed RNA Bind-n-Seq (RBNS) to test the ARID domain’s capability to interact with specific RNA motifs. This *in vitro* high-throughput assay allows to identify the binding preferences of an RBP^43,62^. A pulldown is performed with a 20 nt random RNA pool flanked by short constant adapter sequences with different concentrations of Strep-tagged RBP (ARID_49-152_: 0.25, 1, 5 µM). The constant regions are then used to add sequencing adapters for subsequent analysis by next-generation sequencing. We obtained ∼35-50 million unique reads for each ARID protein concentration. By comparing the frequencies of k-mers in the input library with the pulldown libraries we were able to identify enriched 6-mers (**Fig. 5b and Data Table 1**). The motifs found here can be broadly divided into two types: (i) those that contain AGGC as a core motif and no uracil, and (ii) those that are rich in AU. Analysis of enriched 5- and 7-mers yielded similar results (**Suppl. Fig. 12a** and **Data Table 1**). Complex binding motifs were calculated to get an insight into the environment of the binding sites (**Suppl. Fig. 12c**). Clustering of the AGGC core motifs results in the 9-mer (A)CAGGCA(GG) (**Fig. 5c**). Based on this, we designed two reverse-complementary 9-mer RNAs in agreement with our minimal length for affine DNA-binding (**Fig. 5c and Suppl. Fig. 3**). The structural features of the identified binding motifs were estimated by calculating the average base pairing probability with RNAfold *in silico*. Here, the AGGC-containing motifs show almost no base pairing and appear unstructured, while the AU-rich ones show no particular preference for being structured or unstructured (**Suppl. Fig. 12d**).

We tested ARID_49-152_ binding to the (A)CAGGCA(GG) motif both as ssRNA with the forward (fw) and reverse (rev) strand individually as well as their annealed dsRNA (**Fig. 5c**). Comparison of CSPs similarly to RBNS data reveals a clear preference of ss vs dsRNA; yet within the ssRNA context, specificity for a defined motif is not particularly pronounced (**Fig. 5d and e and Suppl. Fig. 12d**). Unexpectedly, the protein regions interacting with ssRNA are the same as are interacting with dsDNA, with residues 80-90 and 120-130 showing the highest CSPs, which raises the question of a so-far unknown single-stranded nucleic acid-binding mode by ARID. Furthermore, titrations with the RBNS-based ssRNAs indeed show stronger CSPs compared with the ADE despite the larger size and the previously suggested specificity of Arid5a for the ADE interaction mediated by the ARID domain^22^. Those findings are in line with the lack of ADE-related motifs observed in RBNS. Altogether, our data suggest a previously unknown RNA sequence preference specific to the core ARID domain, which may indicate possibly unidentified (m)RNAs bound by Arid5a *in vivo*.

### IDRs increase the ARID RNA-binding affinity in a length-dependent manner

To this stage, our data suggest the ARID core domain to prefer ssRNA over dsRNA and folded RNA and an obvious contribution of the C-terminal IDR to the binding affinity for the ADE. Consequently, we wondered how the ARID-extending IDRs would influence the RNA-binding capacity of other RNA sequences that are longer and more complexly folded. We started with a prolonged dsRNA (19mer_ds), with a central AU-rich core and GC-stabilized flanking regions (**Fig. 6 and Suppl. Fig. 13**). We recorded ^1^H-^15^N-HSQCs of the minimal core and the extended ARID domain in presence and absence of RNA. The core ARID domain interacted only weakly with the 19mer_dsRNA, indicated by minor CSPs in the spectral overlay (**Fig. 6a**). In contrast, severe line-broadening of the IDR-extended ARID_37-183_ suggested a strongly increased interaction with the 19mer_dsRNA (**Fig. 6b**). This suggests an increasing contribution of IDRs in the case of extended RNA stretches, likely reasoned by the steric possibilities and high density of charges.

**Fig. 6:**
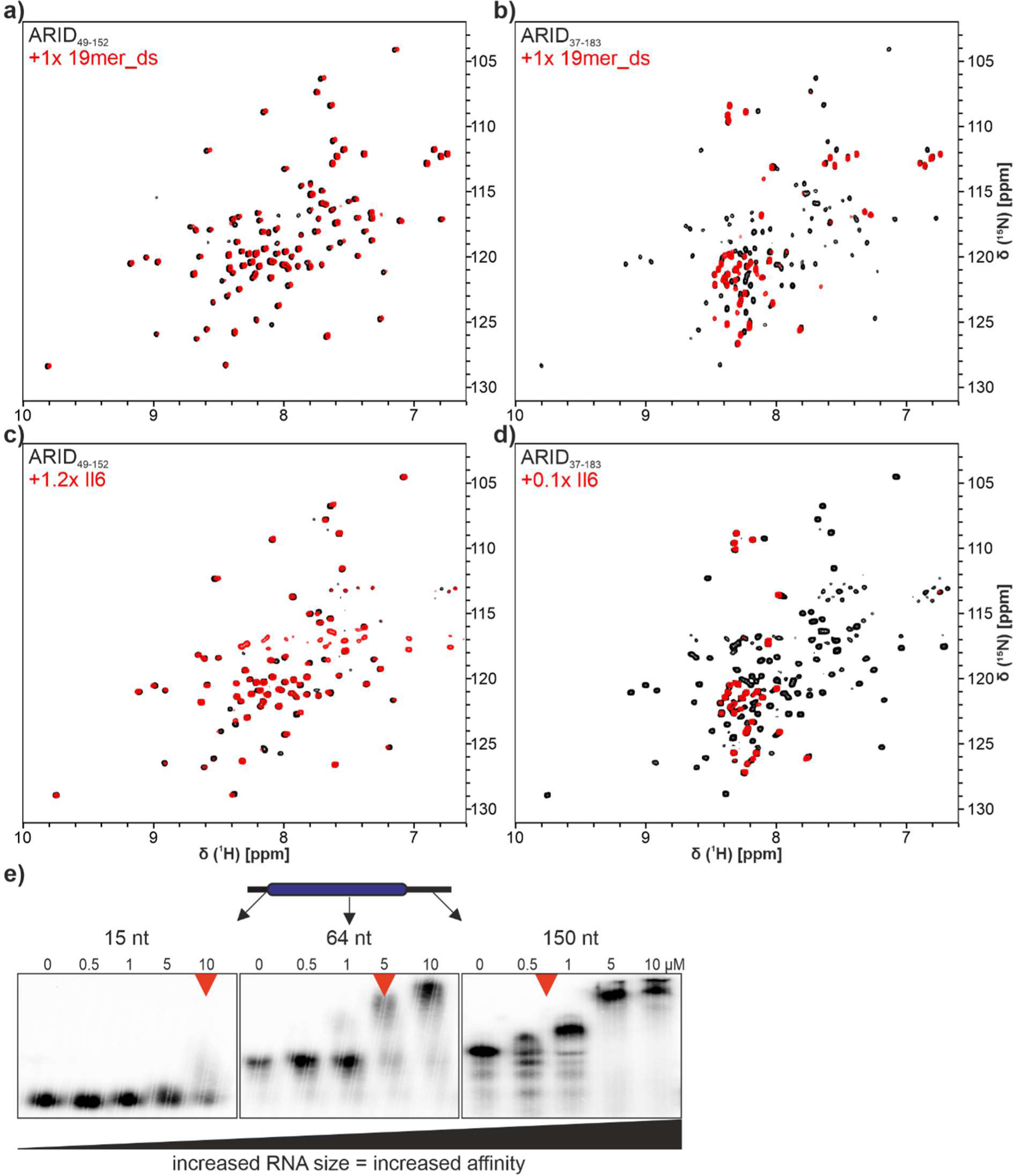
Arid5a ARID domain binding to RNA. **a)**+**b)**: ^1^H-^15^N-HSQCs of ARID_49-152_ (**a**) or _37-183_ (**b**) without RNA (black) or with 1x 19mer_ds RNA (red). **c)**+**d)**: ^1^H-^15^N-TROSY-HSQC of ARID_49-152_ (**c**) or _37-183_ (**d**) without RNA (black) or with 1.2x and 0.1x human Interleukin-6 mRNA (red) respectively. Protein concentration for all NMR measurements was 50 µM. **e)** EMSAs showing that increased RNA size leads to more affine binding by ARID_37-183_. Red arrows indicate approx. 50% bound DNA (as judged by optical inspection).

To test this hypothesis, we used a previously described physiologically relevant target sequence^13^ located in the 3’-UTR of the *Il-6* mRNA. The 129-nt sequence likely represents the naturally occurring RNA-folding complexity, providing stretches of ssRNA (loops) and base-paired regions (**Suppl. Fig. 13g**). Strikingly, while the ARID core domain still binds with only moderate affinity to the RNA in 1.2-fold molar excess, the extended ARID_37-183_ strongly interacts already in sub-stoichiometric concentrations (<0.1x) (**Fig. 6c and d**). Our results suggest that the ARID IDRs drive RNA-binding affinity in dependence of the provided density of negative charge. To confirm this hypothesis, we further compared EMSA-derived apparent affinities to RNAs of increasing length and see a clear correlation between affinity and RNA length (**Fig. 6e**). Likely, this effect is also supported by more than one protein binding to the larger RNAs (see right panel).

In conclusion, our results show that Arid5a is principally capable of interacting with RNA. The intrinsically low affinity of the core ARID domain is compensated by its IDR extensions in a nonspecific manner. Consequently, those nonspecific interactions favor RNAs of increasing size, while the core ARID domain remains restrictive to very specific sequences.

### iCLIP2 reveals that Arid5a binds to ssRNA in cells with a preference for U-rich stretches

Our data so-far suggest that Arid5a is capable of tight interactions with available RNAs, albeit primarily driven by charge interactions. On the other hand, the core ARID domain shows a particular preference for short motifs, which still may steer specific interactions with RNAs more selectively. We speculated that those motifs will be embedded in a larger RNA context *in vivo*, where RNP complex formation will be supported and modulated by the presence of the ARID-flanking regions, and potentially further regions of Arid5a beyond that. Motivated by those assumptions, we performed individual-nucleotide resolution UV crosslinking and immunoprecipitation (iCLIP2)^46^ with full-length Arid5a in murine P19 cells. To our knowledge, this has been the first CLIP experiment carried out with an Arid protein to date.

Murine P19 cells express *Arid5a* mRNA (**Suppl. Fig. 14a**), but a specific antibody suitable for iCLIP is lacking. Hence, we generated an expression plasmid with the murine full-length Arid5a (see sequence alignment and conservation with human Arid5a in **Suppl. Fig. 2**) fused to a C-terminal GFP-tag (mArid5a-GFP, **Suppl. Table 2**). P19 WT cells were transfected in three replicates and subjected to the iCLIP2 procedure using an anti-GFP antibody (**Suppl. Fig. 14b**, see material and methods). All three replicate experiments were highly reproducible and gave rise to more than 7 million crosslinks (**Suppl. Fig. 14b and c**) and 9,895 binding sites with an optimal width of 7 nucleotides (nt) that were used for downstream analysis. Arid5a binding sites are found predominantly in 2,607 protein-coding genes but also in 48 lncRNAs and other noncoding RNAs (**Fig. 7a**). A 3-mer enrichment analysis reveals that Arid5a crosslinks preferentially at U-rich stretches (**Fig. 7c**). The ramp-like enrichment pattern indicates that Arid5a sits at the very 3’-end of polyU stretches (**Fig. 7c and e**). We hypothesized that this positioning of Arid5a may result from a specific interaction downstream of the polyU stretches via the ARID domain, while its adjacent IDRs crosslink to Us in a fixed Arid5a-RNA orientation. U-rich stretches were also found enriched by RBNS (**Suppl. Fig. 12b and c**), suggesting that this preference is not merely due to a UV crosslinking bias for U. To test our hypothesis, we searched for enriched 3-mers downstream of the Arid5a binding sites. Of note, we observe a general enrichment of AG-rich 3-mers (**Fig. 7d**). Moreover, the four 3-mers CAG, AGG, GGC and GCA contained within the RBNS-enriched consensus motif (A)CAGGCA(G) (**Fig. 5b and c**) are consistently enriched downstream of the binding sites (**Fig. 7d and f**), suggesting that Arid5a shows a preference for this RNA motif also *in vivo*. The positioning of these 3-mers downstream to the polyU stretches might help to position the binding of the ARID domain.

**Fig. 7:**
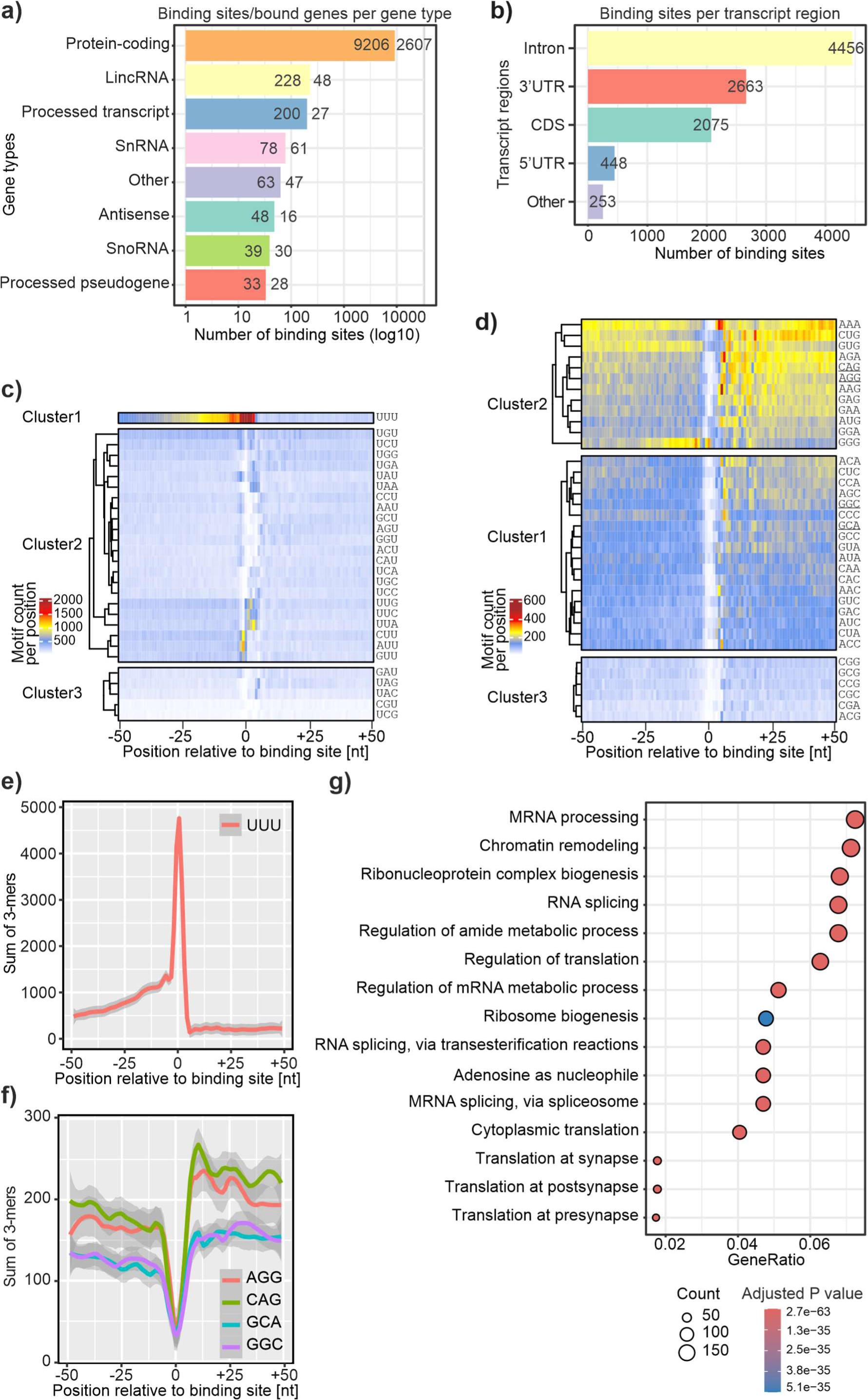
Binding preferences of Arid5a in endogenous RNAs determined by iCLIP2. **a)** Binding site distribution of Arid5a in different gene biotypes. lincRNA, long intergenic non-coding RNA; snRNA, small nuclear RNA; snoRNA, small nucleolar RNA. **b)** Arid5a binding sites in transcript regions of protein-coding genes. UTR, untranslated region. CDS, coding sequence. **c)** Heatmap showing clusters of 3-mers starting/ending with U around Arid5a binding sites in a window of ±50 nt. **d)** Heatmap showing clusters of all other 3-mers around Arid5a binding sites in a window of ±50 nt. RBNS-derived 3-mers are underlined. **e)** Frequency of UUU per position in a window of ±50 nt around the binding sites. **f)** Frequency of the 3-mers AGG, CAG, GCC and AGC from the RBNS consensus motif per position in a window of ±50 nt around the binding sites. **g)** Functional enrichment analysis (Gene Ontology Biological Process) using for transcripts with Arid5a binding sites.

Looking at the bound transcripts, we observe Arid5a across all transcript regions, including introns, 3’UTRs and exons (**Fig. 7b**), indicating that Arid5a binds to pre-mRNAs in the nucleus. Arid5a targets are enriched for transcripts involved in mRNA processing, chromatin remodelling and translation regulation (**Fig. 7g**). Altogether, our data suggest that Arid5a is a sequence-specific dual DNA- and RNA-binding protein that binds to a subset of transcripts *in vivo* and may have an accessory function in chromatin-related transcript processing, or in transcription regulation, possibly in an RNA-supported context (see discussion).

### Arid5a is a strictly nuclear protein and co-localizes with heterochromatin

The proposed dual function of Arid5a in gene regulation both via interaction with DNA/chromatin and protection against mRNA degradation in the cytoplasm^13,21,22^ requires the protein to be present in both the nucleus and the cytoplasm. However, our iCLIP2 data show that Arid5a binds preferentially to unspliced pre-mRNAs, suggesting an exclusively nuclear function of Arid5a associated with chromatin. To test the subcellular localization of Arid5a under normal conditions, we performed confocal fluorescence microscopy of P19 wild type (WT) cells transfected with mArid5a-GFP. As controls we transfected SRSF3-GFP as marker for the nucleoplasm^63^ and performed immunofluorescence (IF) for G3BP1 as a cytoplasmic marker. A plasmid expressing GFP alone was used as an ubiquitously present protein^64^.

Arid5a clearly localizes to the nucleus, and no signal is detectable in the cytoplasm (**Fig. 8**). However, Arid5a shows a markedly distinct localization pattern compared to the splicing regulator SRSF3, which is found in nuclear speckles and the nucleoplasm. Arid5a perfectly co-localizes with some of the bright heterochromatin dots, indicating a close proximity to silenced chromatin. Together with our iCLIP2 data this suggests that Arid5a might use its dual nucleic acid-binding capability to interact with DNA and pre-mRNA simultaneously, e.g. to detect transcribed loci and then modulate transcriptional repression as it was described earlier^17,18^ and very recently for TF with an RBP activity^65^.

**Fig. 8:**
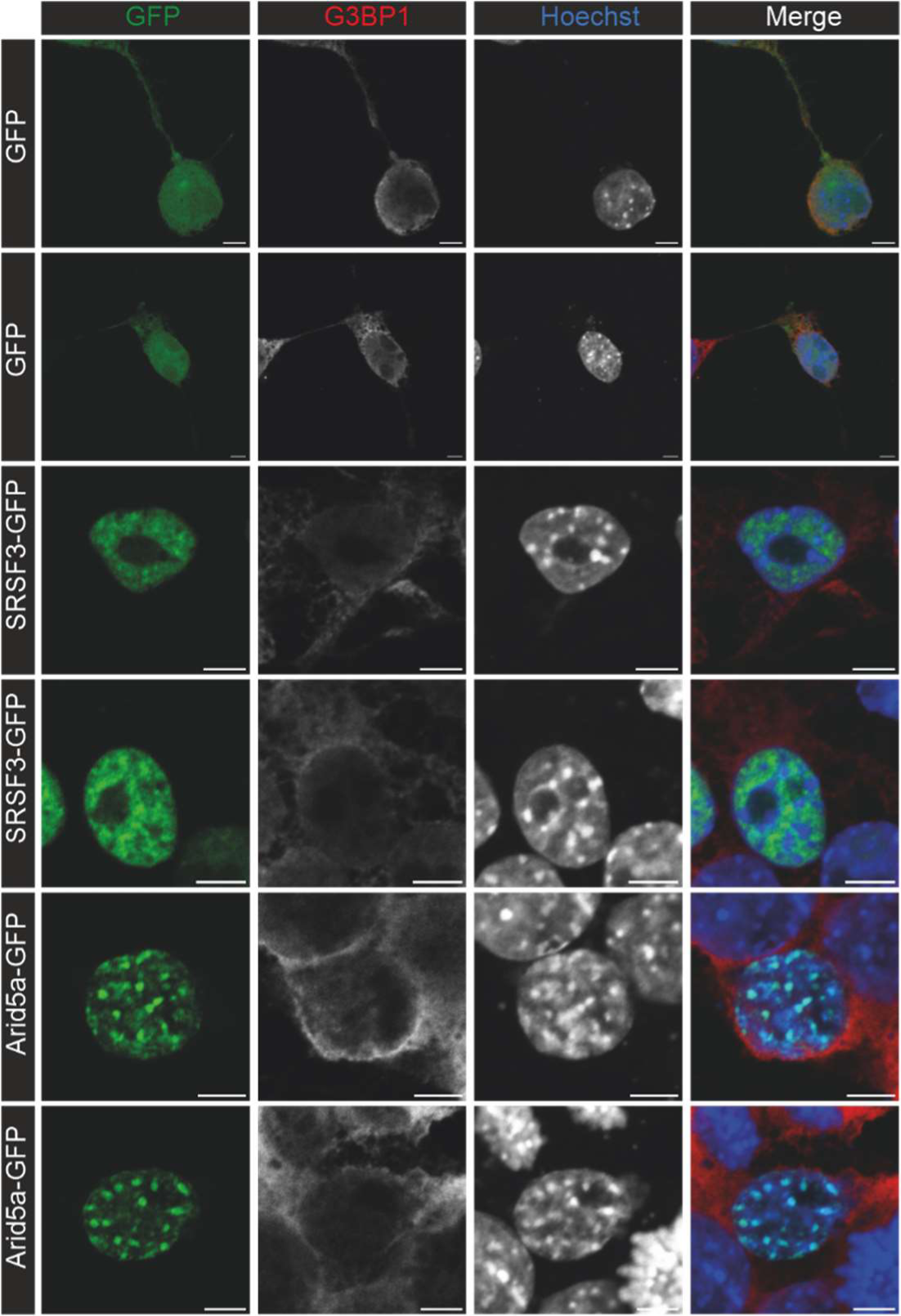
Subcellular localization and RNA-binding of Arid5a. Representative confocal micrograph showing that Arid5a-GFP localizes to the nucleus of murine P19 cells and co-localizes with bright heterochromatin dots. GFP was used to stain the entire cell and SRSF3-GFP as marker for nuclear speckles. Staining for G3BP1 in all cells labels the cytoplasm, and Hoechst stains chromatin. All zoom-ins are twofold. Scale bars = 5 µm.

### RNA-binding is a more widespread capacity of ARID domains

Prior to this study, RNA-binding had only been described for Arid5a, but for none of the other 14 human Arids. Driven by the observations for Arid5a above, we wondered if RNA-binding was a more general feature within this protein family. To test this, we used the closely related Arid5b as well as Arid1a and Jarid1a to perform analogous ^1^H-^15^N-HSQC measurements with and without the 19mer_dsRNA (**Suppl. Fig. 13a-e**). Surprisingly, all three Arids showed obvious line broadening with Jarid1a being most affected and very similar to ARID_37-183_. Arid5b and Arid1a showed less, but still mentionable line broadening. This indicates that also other Arids are able to bind RNA *in vitro* and hints at so-far unexplored functions of these proteins. To corroborate protein-observed HSQC spectra, we recorded imino proton spectra of the 19mer_dsRNA with and without the Arids in order to examine the influence of protein binding on the RNA chemical shifts (**Suppl. Fig. 13f**). We did not observe significant CSPs for the imino peaks, but minor line broadening indicates that all Arid proteins interact with the RNA backbone, suggesting little specificity for the herein provided RNA motif. Nonetheless, these results reveal that some – if not all – Arid proteins are generally able to interact with RNA through their respective ARID domains and RNA-binding competence is thus not a unique observation for Arid5a. This opens up interesting questions for further detailed studies in the future into whether and how RNA-binding is of functional relevance for them (as suggested for Arid5a).

## Discussion

A recent study suggests more than 100 TFs are actively involved in splicing through their DRBP function^66^. The ability to interact with both DNA and RNA is either conferred by a combination of specialized domains, e.g. in Sox2^10^ or SAFB2^67^ or by the dual exploitation of one domain^12,68^. Recently, the role of IDRs for DNA- and RNA-recognition, often via the same sequences^69^, has come into focus, but specificity parameters like in Arid5a remain elusive based on the lack of structure-derivable knowledge.

Arid proteins are categorized as exclusive DNA-binders with only one exception: Arid5a is capable of binding RNA, with specific target mRNAs and a responsible folded motif presented in earlier work^2,13^. However, not only the structural basis of this unique observation has remained unresolved, but also a clear understanding of the precise target nucleic acid preferences of Arid5a, all of which are expected to involve regions beyond the core ARID domain. In support, prior data on Arid5a RNA-binding had been achieved with the full-length protein^21,22,28^, while RNA-binding is abolished in the absence of the ARID domain^13^. The latter, as well as a study involving a mutant within the core ARID^21^, claim that the domain is sufficient for RNA-binding, but ignore contributions from sequence elements directly adjacent. Altogether, an atom-resolved proof of the Arid5a ARID domain interacting with DNA and RNA in an isolated, *in vitro* setup had been missing.

We here provide a detailed interrogation of the Arid5a ARID preferred DNA target DNA motif, focusing on the core domain, but taking into account contributions of N- and C-terminally extending IDRs. With a core “AATA/TATT” sequence, we find that the core ARID prefers a similar DNA target motif as its related family partner Arid5b^15,16^. This was unexpected considering the core ARIDs of both proteins share a sequence identity of only 70.2 %, and the extended domains an even lower 58.3 %, respectively. Interestingly, the regions involved in DNA-binding (L1 and H4-L2-H5) share a sequence similarity of 97.6% and identity of 85.4%, explaining their preference for identical DNA motifs (**Fig. 1b**). This is further supported by the finding that amino acids analogous to L2-residue T125 in Arid5a are determinants of DNA preference^56^. T125 is both conserved in Arid5a between species, and between Arid5a and 5b. Finally, early work had already suggested Arid5a to interact with multiple AT-rich sites, but not with a precise motif^17^. In contrast, other members of the Arid family do not necessarily prefer AT-rich sequences, as e.g. reported for Arid1a^55,58^ and JARID1a^56^ with a serine and lysine, respectively, at this position. Our data (**Fig. 2**) confirm that Arid1a and JARID1a can bind AT- and GC-rich DNA equally strong. Similarly, we do not confirm Jarid1a to exclusively bind GC-rich DNA, thus contradicting the previous suggestions^56^.

The co-existence of Arid5a and 5b in higher eukaryotes remains enigmatic, seeing their shared DNA target motif preference of the core ARID domain. Notably, literature does not list an overlap of genes regulated in transcription. Our findings suggest that a fine-tuning of DNA targets may take place through modulation by the non-identical IDRs. This is supported by earlier findings, in which Arid5b was shown to interact with DNA via its C-terminal extension^61^.

Our data show that the positively charged C-terminal IDR also supports affinity of Arid5a to DNA. Notably, the NMR data reveal a larger relative contribution to binding of GC DNA. This suggests this region provides a general support in DNA engagement, ultimately allowing the core ARID domain to selectively encounter AT motifs (**Fig. 9a**). While here, we suggest opposing charges to drive encounter complex formation, a recent study found negatively charged IDRs to accelerate specific motif search^70^, likely preventing too tight interactions. We, however, did not find a similar contribution from the negatively charged N-terminal extension. Certainly, nature has established multiple modes of fine-tuning DNA recognition through IDRs^71-73^, including roles for hydrophobic sequences as recently shown by Jonas et al.^74^.

**Fig. 9:**
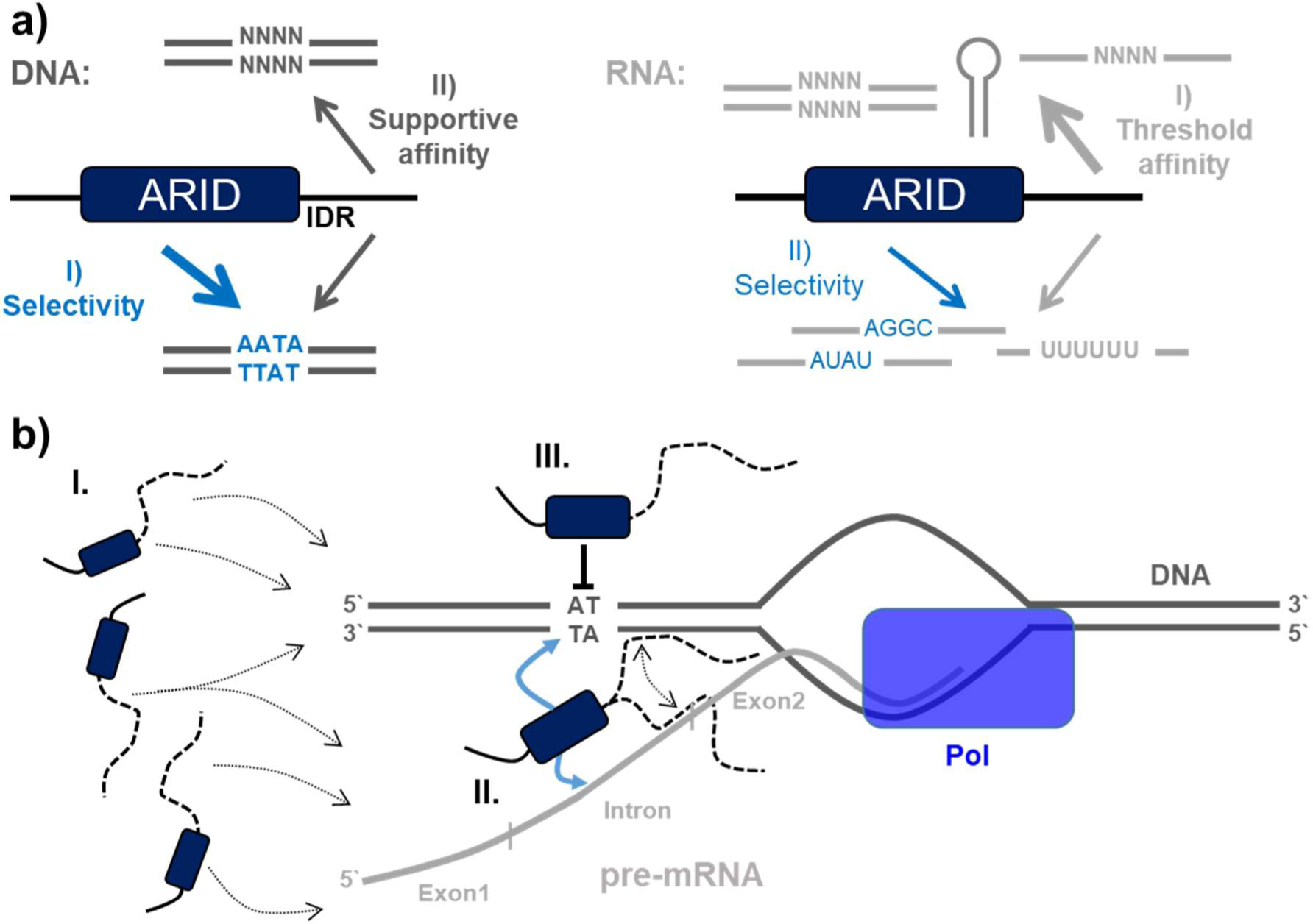
Summary of Arid5a interacting with nucleic acids. **a)** Overview of possible and preferred interactions of the extended ARID domain, ARID_37-183_, with DNAs (left) and RNAs (right) with an apparent hierarchy of affinity and selectivity. **b)** Hypothetical model of transcription modulation by Arid5a integrating the in vitro and in vivo findings of Arid5a’s specific and non-specific nucleic acid-interactions mediated by the core domain (dark blue rectangle) and the IDR extensions (broken lines): Recruitment of Arid5a to DNA/RNA (‘scanning’) will primarily locate the protein to AT-rich DNA promoter/enhancer regions (I). Increase of local Arid5a concentration through recognition of intron-exon boundaries in nascent transcripts closely located to transcribing DNA (II). Arid5a binding could allow productive transcription simply by relieving the block of DNA promoter/enhancer regions. Alternatively, Arid5a recruitment to pre-mRNA can lead to recognition of the gene’s cognate promoter region and thus, its silencing.

The strong similarity in DNA-binding between Arid5a and Arid5b raises the question why to date only Arid5a was found to bind RNA. A sequence conservation within the extended ARID domains of the two proteins below 60% supports the hypothesis that the IDRs play a central role in (distinctive) RNA-binding competence. For Arid5a, we here unambiguously provide an atom-resolved proof for its proposed dual nucleic acid-binding competence (**Fig. 9a**), while no work had shown RNA-binding by robust *in vitro* experiments before. Unexpectedly, we found only weak binding of the utilized Arid5a constructs to the previously described *Ox40* ADE motif^22-24^, while visibly enhanced by the IDRs. Our RBNS approach (unprecedented for an ARID domain) suggested short ssRNAs, superior to the ADE. In support, these motifs were partially found *in vivo*, demonstrated by the first iCLIP2 experiment with an Arid protein.

In general, binding to sequence- and size-equivalents of DNA (**Suppl. Fig. 15**) revealed the subordinated affinity of the ARID domain to RNA. In fact, we find that RNA-binding shows a CSP pattern reminiscent of non-specific DNA-binding by NMR (**Suppl. Fig. 16**). We do not rule out that we missed a complex-folded RNA motif preferentially bound by the ARID domain, similar to the unique binding of ADE and CDE elements by the Roquin ROQ domain^23,75^.

The nuclear localization of Arid5a is supported by early characterization of the protein in different tissue types^17^. More recent work identified Arid5a as specific RBP in stimulated immune cells, including export to the cytosol^13,21,22,28,76^. Our data reveal a full nuclear localization, while we did not perform iCLIP2 under differential conditions and do not question a possible engagement with specific transcripts outside the nucleus. Still, we doubt Arid5a is a broadly acting RBP, neither in the nucleus nor cytoplasm, as it crosslinked much less to RNA than e.g. the splicing factor SRSF5 with ∼40 times more binding sites in a similar approach^77^. Apart from the above, the general capability of interacting with RNA had not yet been tested for other Arid proteins to our best of knowledge. When we compare findings for Arid5a to Arid1a, 5b, and Jarid1a, our data hint at a more common RNA-binding capability of ARID domains than expected. This suggests unknown functions for Arids with respect to gene regulation, e.g. at the interface of transcriptional and post-transcriptional levels as very recently shown for Arid1a^78^. Considering a difference in affinity between DNA and RNA, as found here for the Arid5a ARID domain, we can speculate whether RNA-binding functions require high protein or target RNA concentrations. In such a scenario, e.g. Arid5a will automatically expand from DNA-binding (transcription regulation) to RNA (transcript)-binding as a consequence of its own abundance. E.g., Arid5a may first act as transcriptional repressor on the chromatin level, while it then stabilizes or blocks transcripts from translation at a later stage, including its abundance-based co-export from the nucleus, thus fulfilling a regulatory role on multiple levels.

In fact, TFs possibly involve simultaneous RNA-binding as a feedback mechanism in transcription, or for recruitment to transcriptional start sites, e.g. via (l)ncRNAs. Similar to the emerging role of circRNAs for RBPs^79^, RNAs may also act as sponges for excessive DBPs^80^ via IDR interactions. DNA- and RNA-binding is a strong indicator for sub-compartmental clustering of transcriptional processes, e.g. for co-transcriptional splicing^66^ or miRNA processing. The latter was suggested for SAFB2^81,82^ as a *bona fide* example of a DRBP^67^. Our iCLIP2 and microscopy data now suggest a similar role for Arid5a, which could function in a mechanism of RNA-induced transcriptional silencing or activation in line with differential regulation of transcription in Arid5a k.o. conditions^83^. Both scenarios will involve the core ARID domain binding to dsDNA and to particular ssRNAs. The observed ramping effect in our iCLIP2 data suggests that Arid5a recognizes steep changes in nucleotide composition, i.e. from longer U-rich stretches to purine-rich sequences (in accordance with RBNS), using its core ARID domain and the flanking IDRs. Such changes in nucleotide composition occur at intron-exon junctions in pre-mRNAs (**Fig. 9b**). We speculate that Arid5a normally binds to DNA, but hops on nascent RNAs in the vicinity when such boundaries emerge and could thereby discriminate normal pre-mRNAs from spurious transcripts. The fact that we detect bound transcripts by iCLIP2 suggests that Arid5a binding rather prevents silencing of the locus, perhaps through loss of DNA-binding, but this requires further investigation.

The support by its adjacent IDRs additionally allows for protein-regulatory features steerable via post-translational modifications. Likewise, IDRs are by default susceptible to proteolysis, an excellent tool to disrupt functional protein moieties^84,85^. Similar to the described PTMs in more distal parts of Arid5a^86^, PTMs in the extended ARID domain may be relevant with respect to DNA vs. RNA preference and general affinity.

We here focused on solution NMR spectroscopy as a valuable tool to *en-detail* correlate chemical shift information with binding modes. CSP patterns, i.e. trajectories, magnitudes and exchange regimes, are unambiguous indicators of a protein domain’s preference for nucleic acid as recently shown in similar studies by us^37,67^ and others^87,88^. As such, CSPs can be used to compare DNA and RNA-binding by Arid5a and consequently the approach is transferable to other nucleic acid-binding domains of interest. The straightforward NMR-centered biochemical setting will on the longer run also allow to unambiguously read-out selective inhibition of one or both DNA and RNA functions as intended for Arid5a earlier^89^.

## Data availability

All sequencing data are available in the Gene Expression Omnibus (GEO) under the accession numbers GSE256029 (RNBS) and GSE254818 (iCLIP2). The tokens for anonymous reviewer access are: axktyqweftgrvkf (RBNS) and ozshmgmyrboldwb (iCLIP2).

## Acknowledgement

We acknowledge excellent technical support by Katharina Targaczewski. We thank Mirko Brüggemann for advice and supervision, and Anke Busch and IMB Bioinformatics Core Facility for processing the iCLIP2 data. Support by the IMB Genomics Core Facility and the use of its NextSeq 500 (funded by the Deutsche Forschungsgemeinschaft [DFG, German Research Foundation] – 329045328) is gratefully acknowledged.

## Author contributions

J.v.E., S.M.K., M.M.M and A.S. designed experiments. J.v.E., S.M.K. and L.I.S. performed NMR- and EMSA-related protein construct cloning, protein production and all corresponding experiments including their analysis. L.O. performed the RBNS assay and its analysis. J.v.E. and F.M. performed iCLIP2 experiments. E.Y. and K.Z. performed iCLIP2 analysis. N.B. transfected P19 cells and took microscope images and provided Arid5a expression data from P19 cells. J.v.E., S.M.K., L.O., J.E.W., K.Z., M.M.M. and A.S. wrote the manuscript.

## Supplemental material

**Supplementary Table 1.**
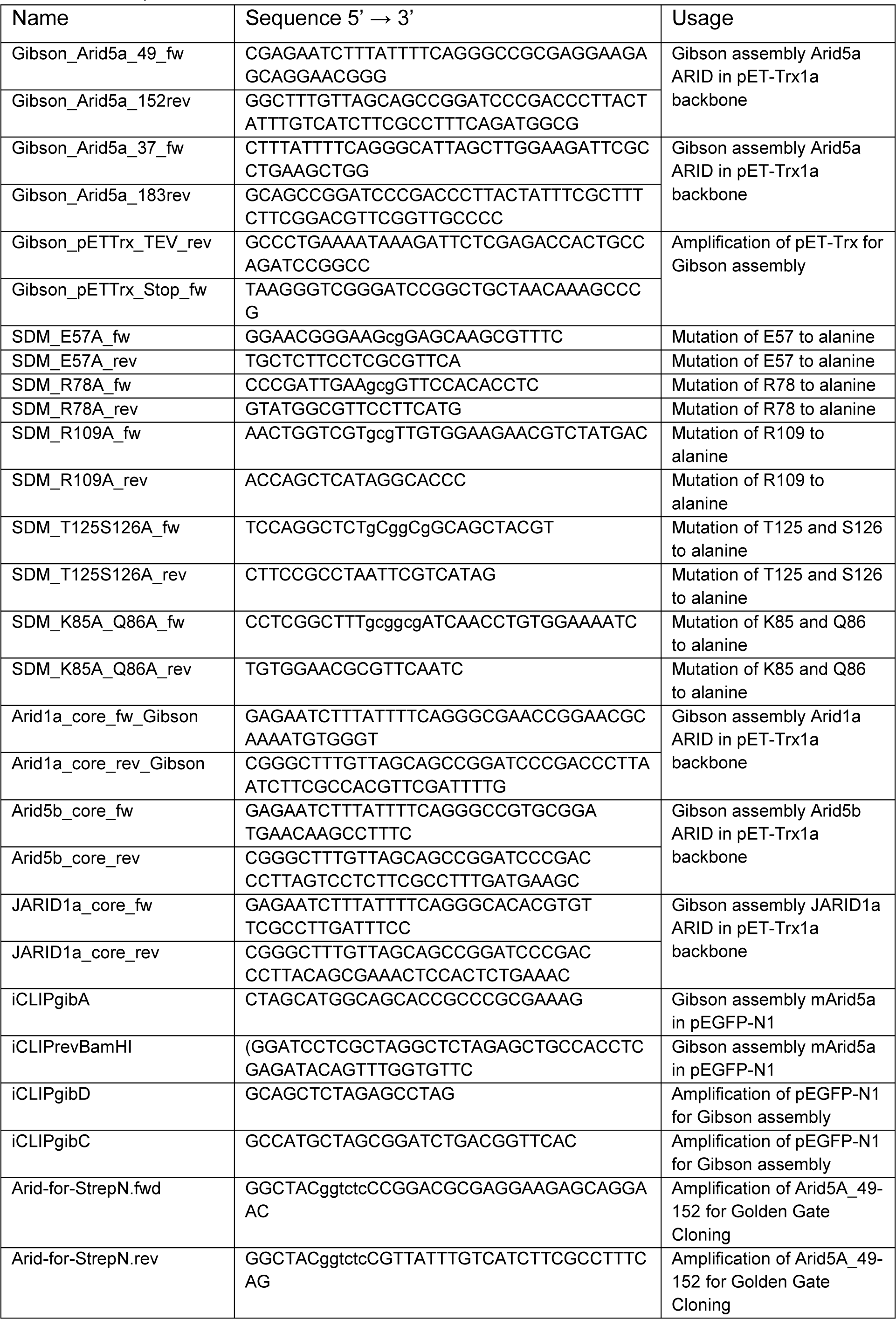
Overview of DNA oligonucleotides used to generate mutants in the ARID domain. Non-capital letters indicate site of mutation.

**Supplementary Table 2.** Vectors used in this study.

| Name | Feature/Background etc. | Application |
| --- | --- | --- |
| pET-Trx | N-terminal His<sub>6</sub>-tag, Thioredoxin-tag, TEV cleavage site, Kanamycin resistance, *lac* repressor, T7 promotor | Cloning and expression of ARID constructs in *E. coli* |
| pEGFP-N1 | N-terminal GFP tag, Neomycin/Kanamycin resistance, CMV promotor | Expression of GFP-tagged murine fl-Arid5a in P19 cells |
| sPHDV64 | Ampicillin resistance, T7 promotor, 3'-HDV ribozyme | *In vitro* transcription of RNAs |
| pET_TRX_Bsa_StrepTag-N | N-terminal His<sub>6</sub>-tag, Thioredoxin-tag, TEV cleavage site, Twin-Strep tag, Kanamycin resistance, *lac* repressor, T7 promotor | Cloning and expression of RBNS-ARID construct in *E. coli* |

**Supplementary Table 3:**
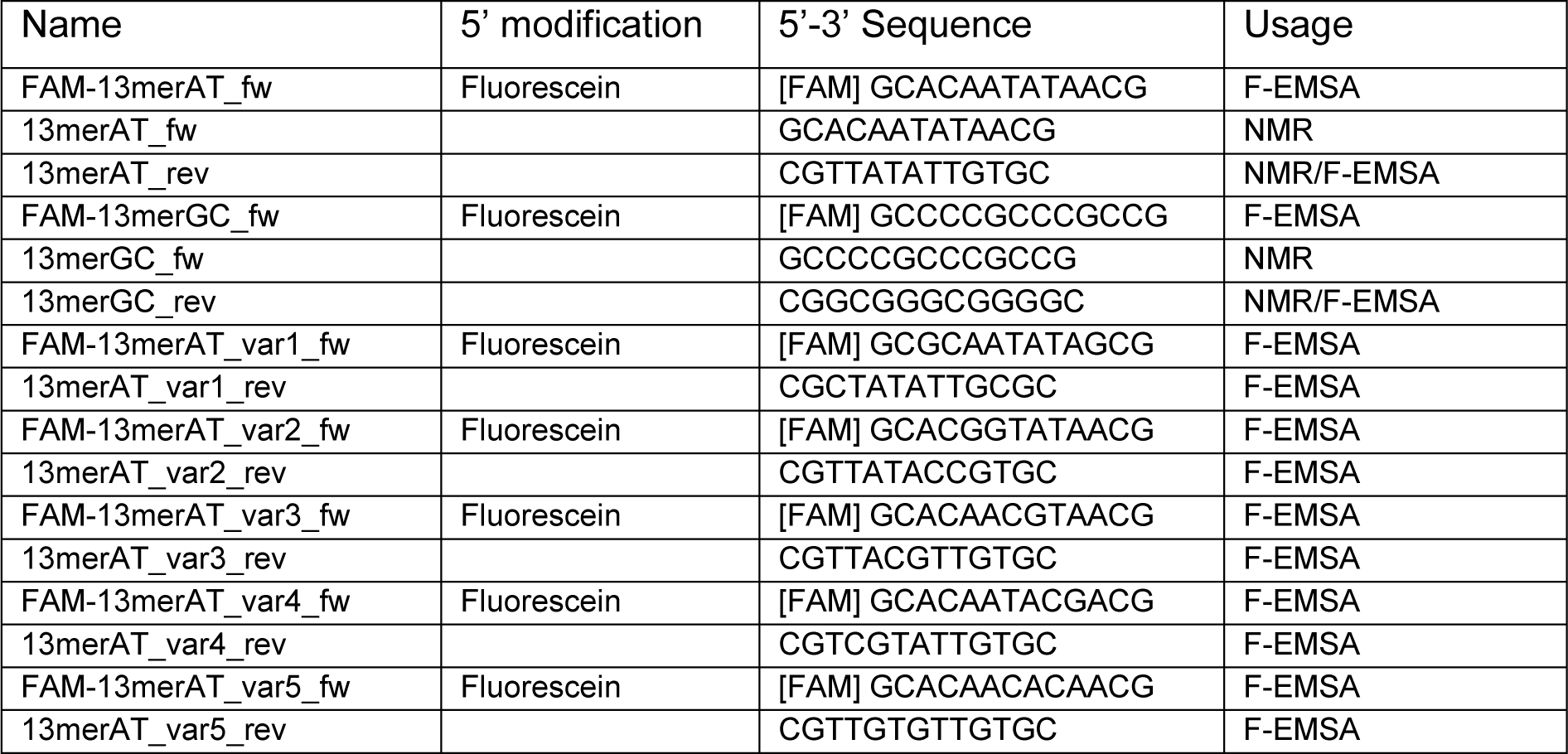
Overview of DNA oligonucleotides used for binding studies with ARIDs.

**Supplementary Table 4.**
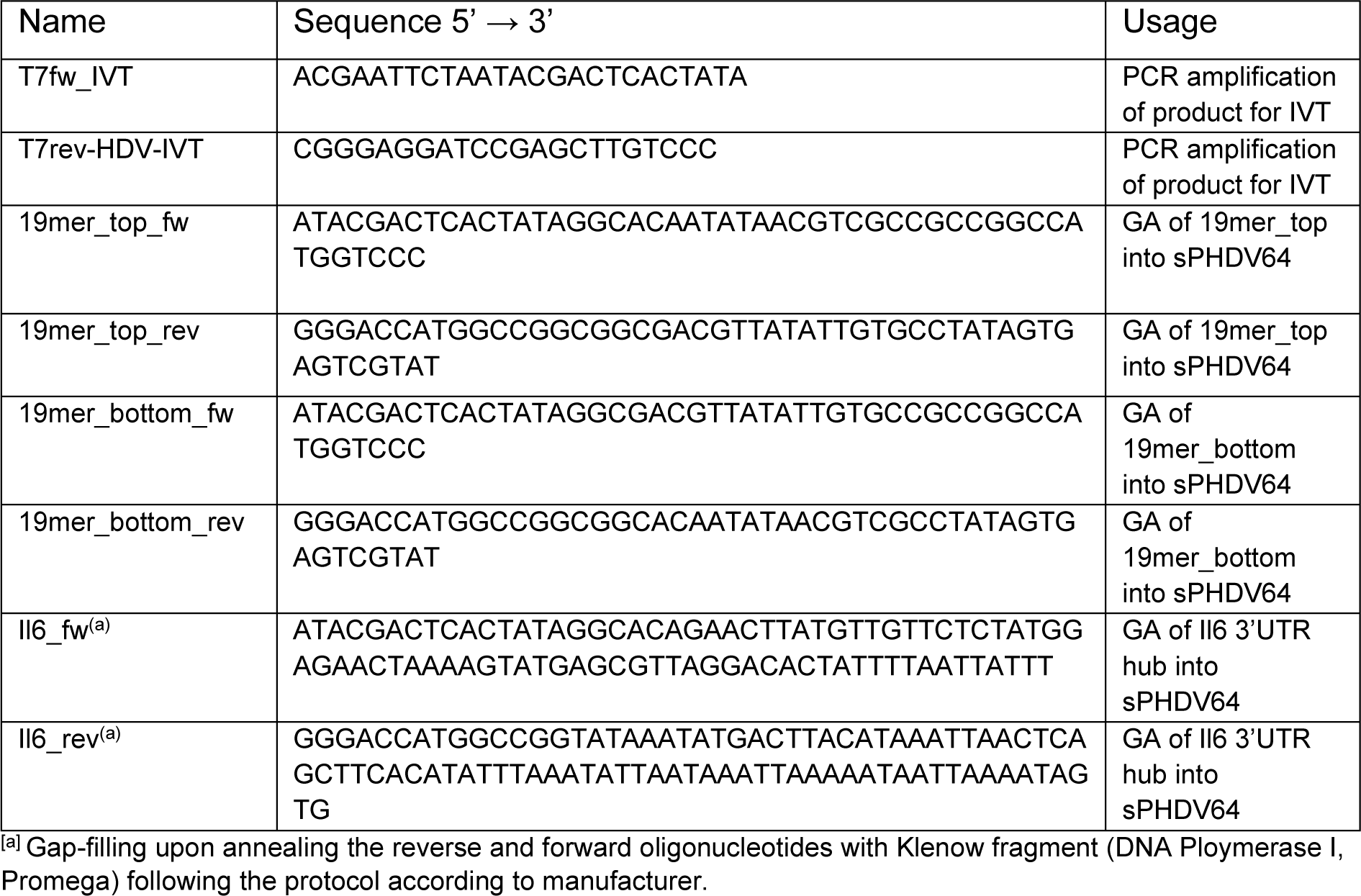
Overview of oligonucleotides for generating IVT templates.

**Supplementary Table 5.**
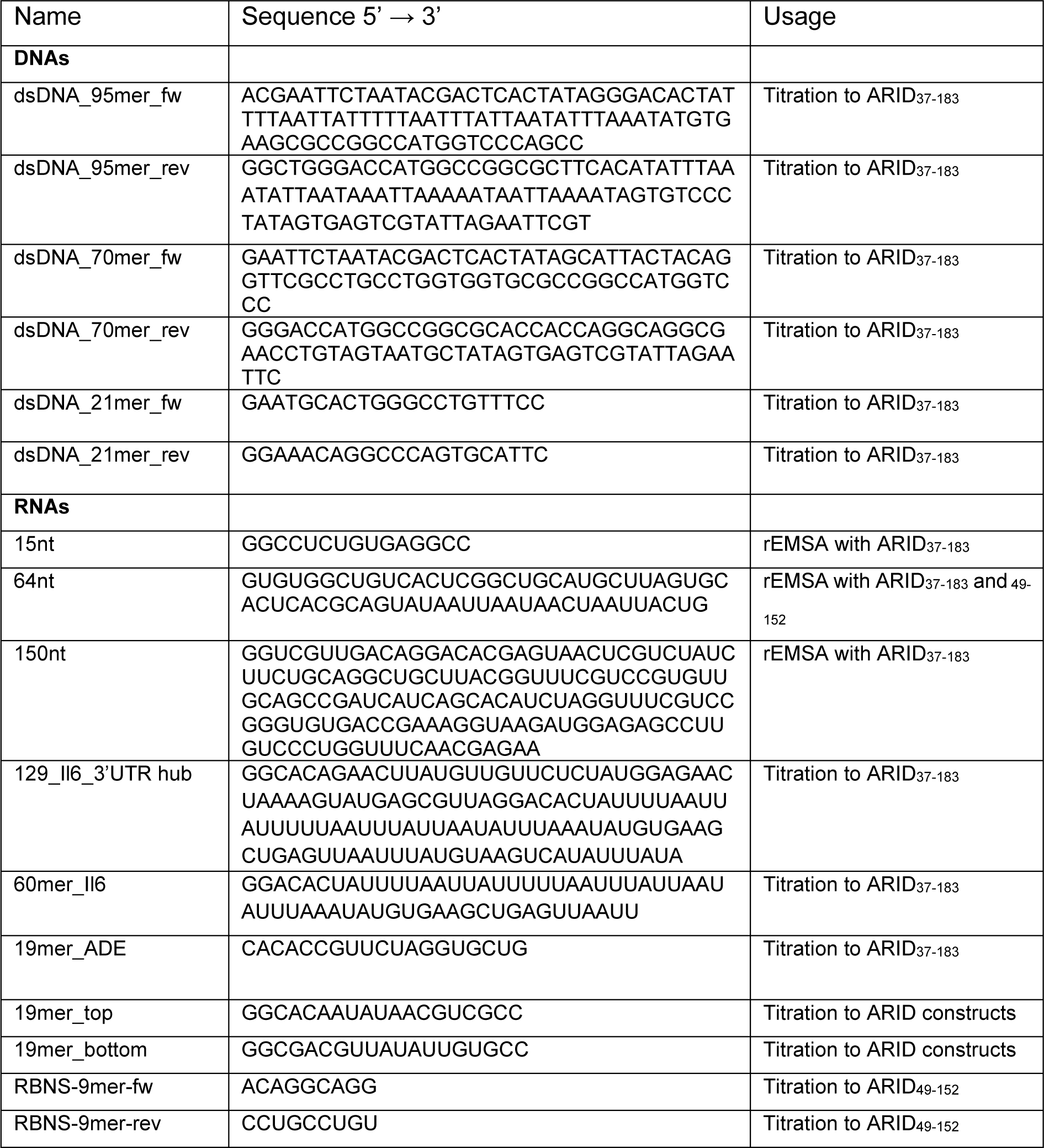
Overview of additional NA oligonucleotides as templates for ARID proteins used in this study. All oligonucleotides were used as non-labelled variants or ^32^P-labelled in house.

**Supplementary Table 6.**
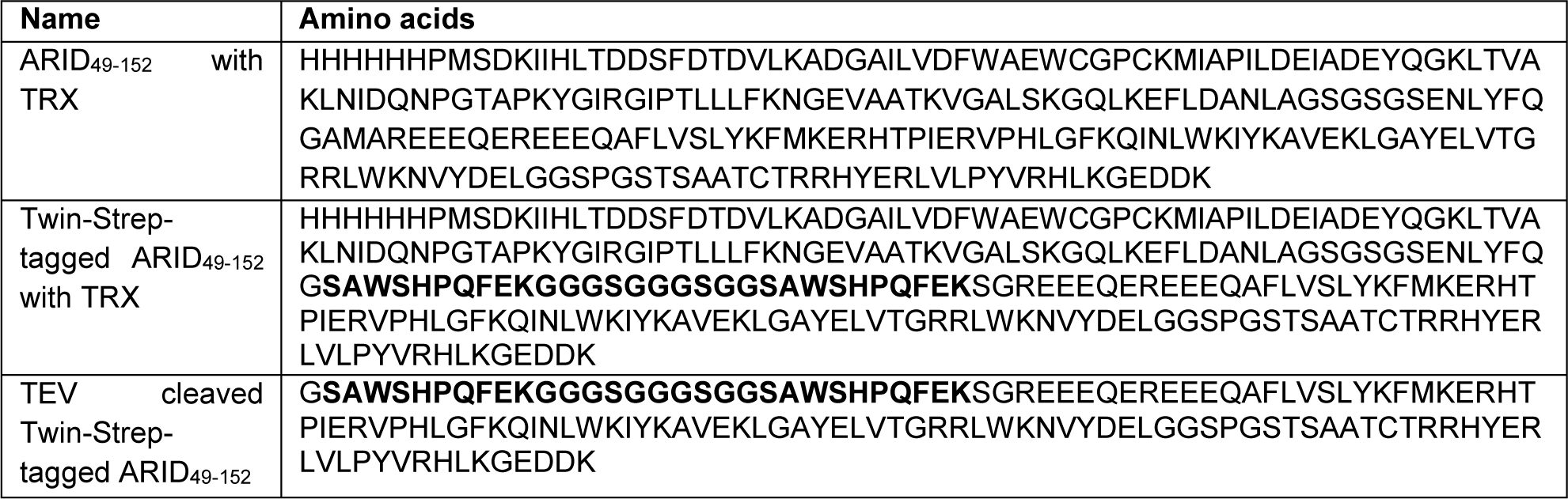
Amino acid sequences of the proteins used for RBNS assay. The Twin-Strep-tag is marked in bold.

**Supplementary Table 7.**
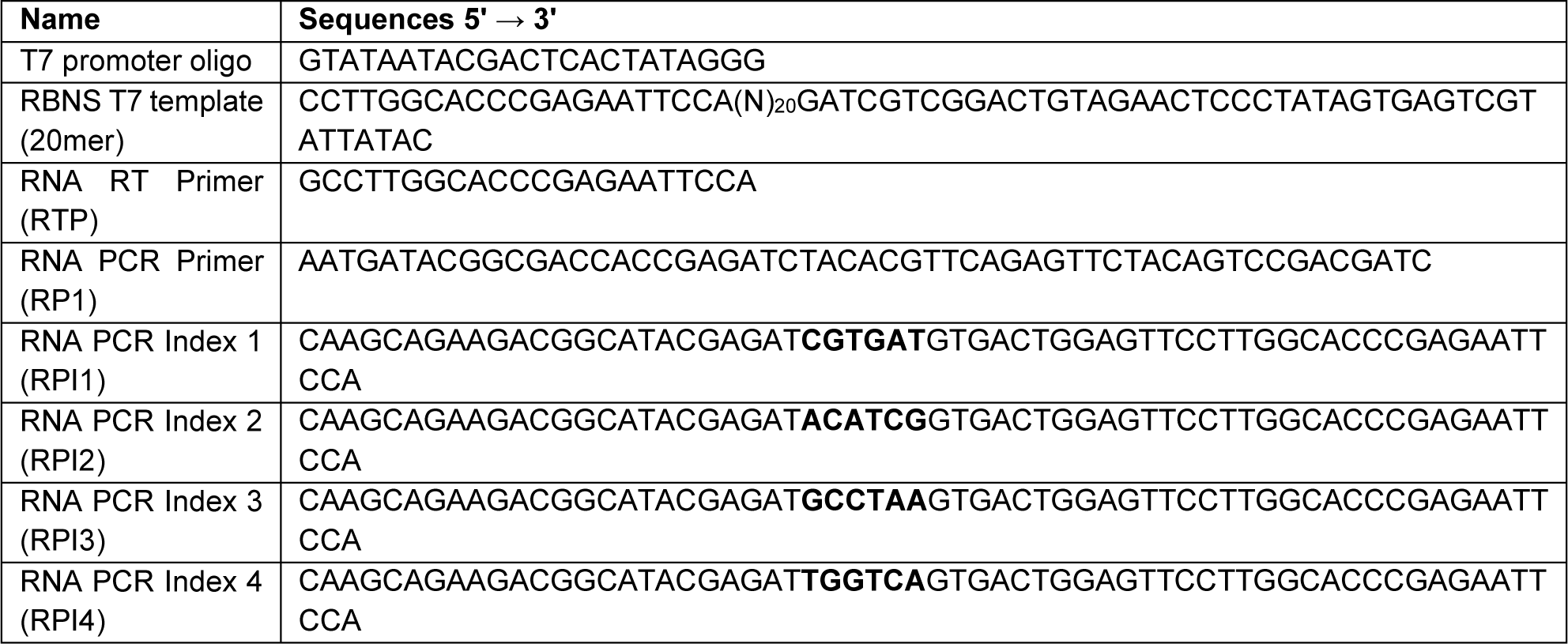
Oligonucleotides used for the RBNS assay.

**Supplementary Table 8.** Indices used for Illumina sequencing.

| Sample Name | i7 Index Name | i7 Index Sequence |
| --- | --- | --- |
| input | RPI1 | ATCACG |
| ARID-0.25-μM | RPI2 | CGATGT |
| ARID-1-μM | RPI3 | TTAGGC |
| ARID-5-μM | RPI4 | TGACCA |

**Supplementary Table 9.**
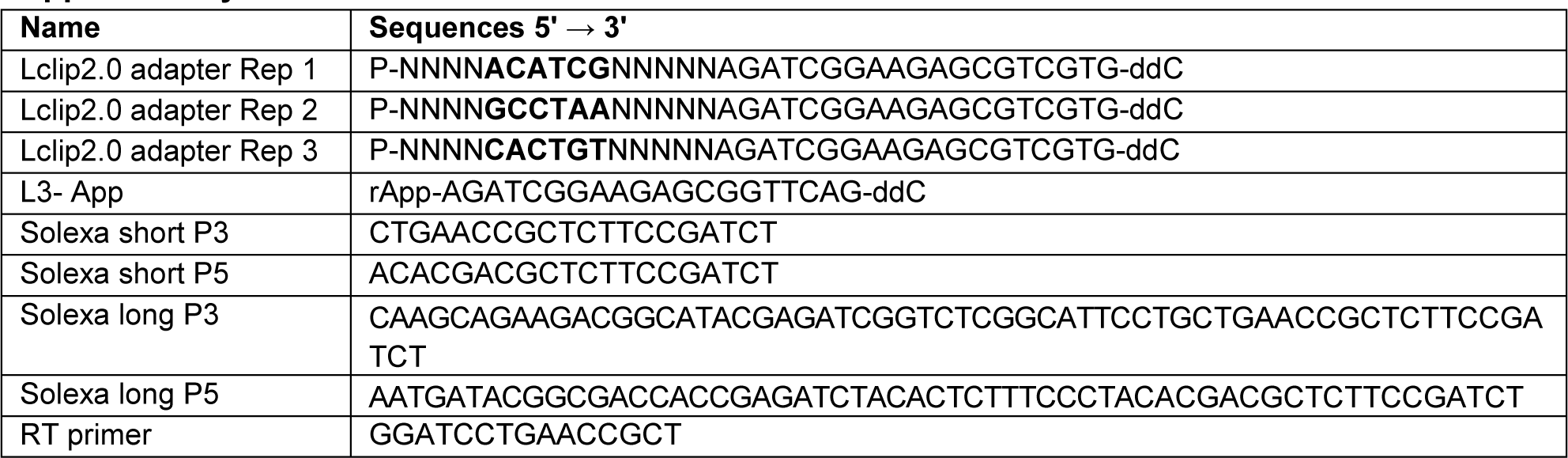
Primers and indices used for iCLIP2.

**Supplementary Fig. 1.**
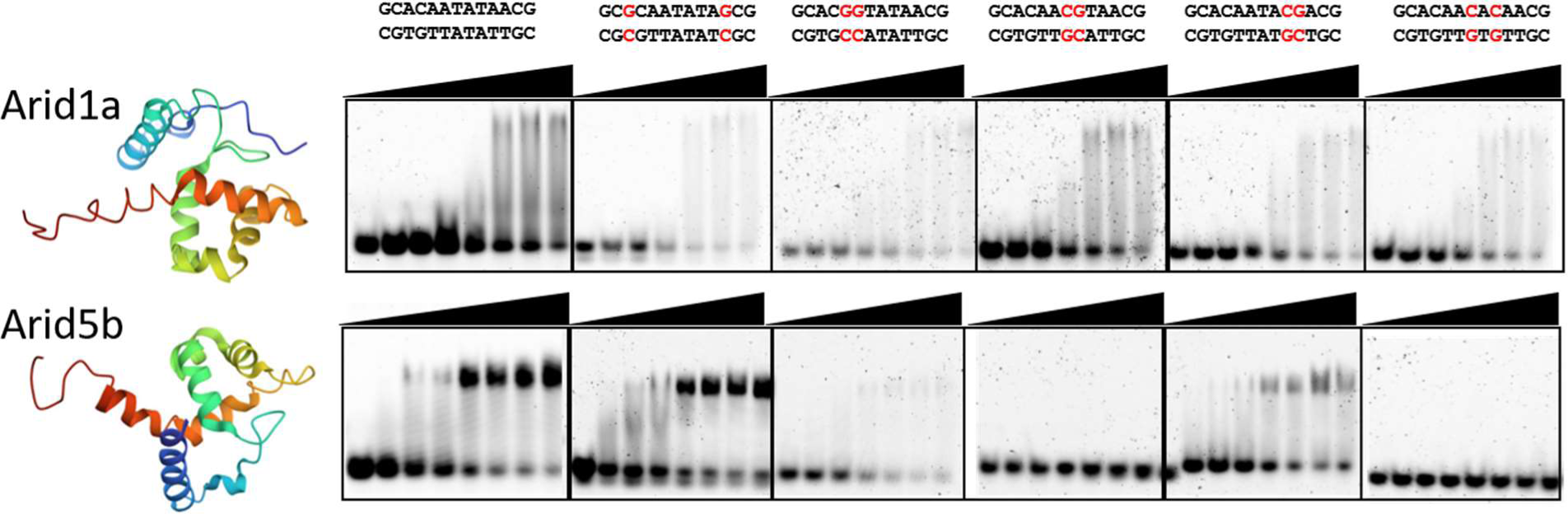
DNA-preferences of selected ARID domains. EMSAs of fluorescently labelled DNAs as given above when titrated with increasing amounts of extended ARID domains from either Arid1a (top) or Arid5b (bottom).

**Supplementary Fig. 2.**
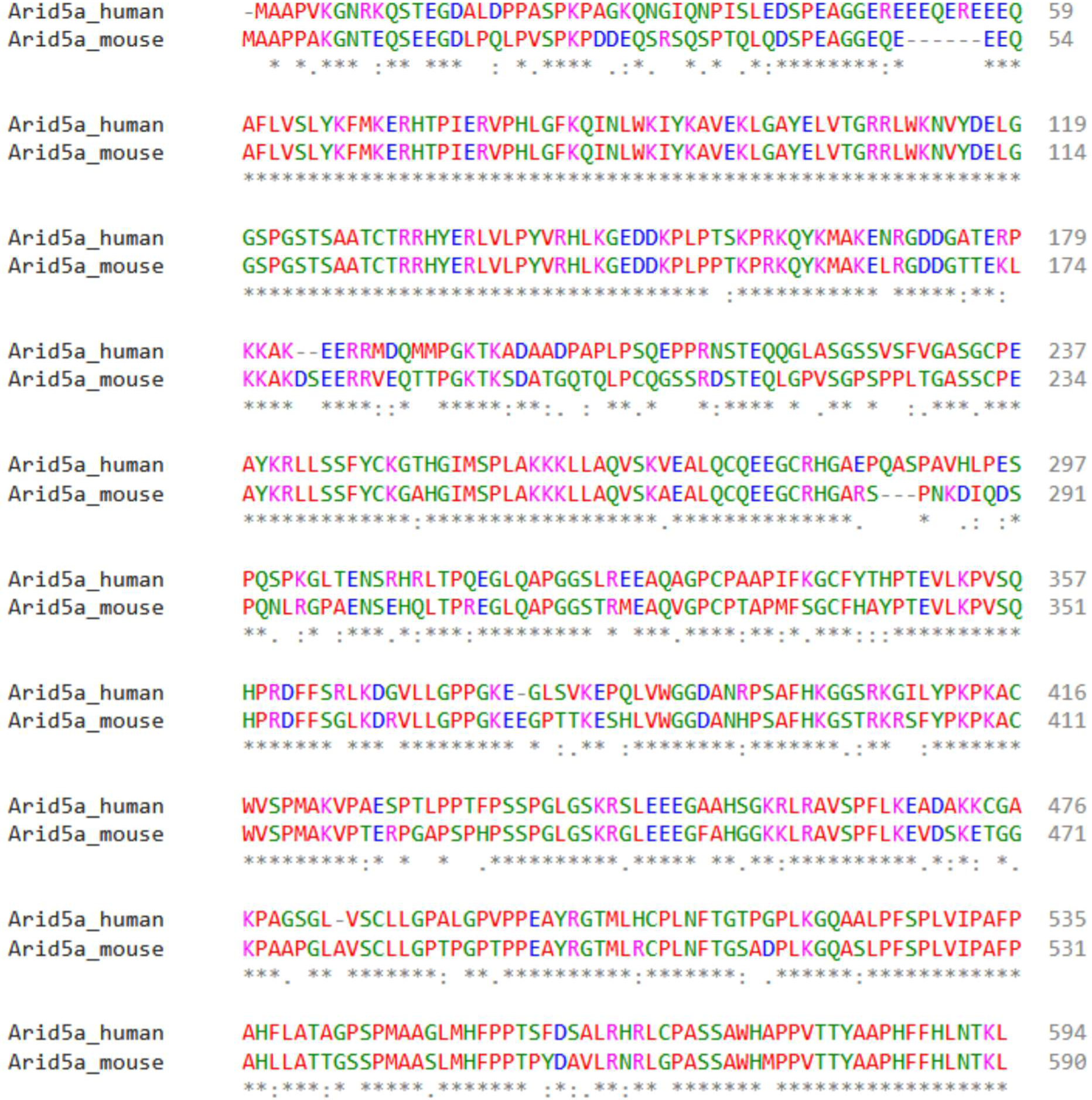
Comparison of complete human and mouse Arid5a protein sequences based on UniProt accession codes Q03989 and Q3U108, respectively. Alignment created with Clustal Omega^1^.

**Supplementary Fig. 3.**
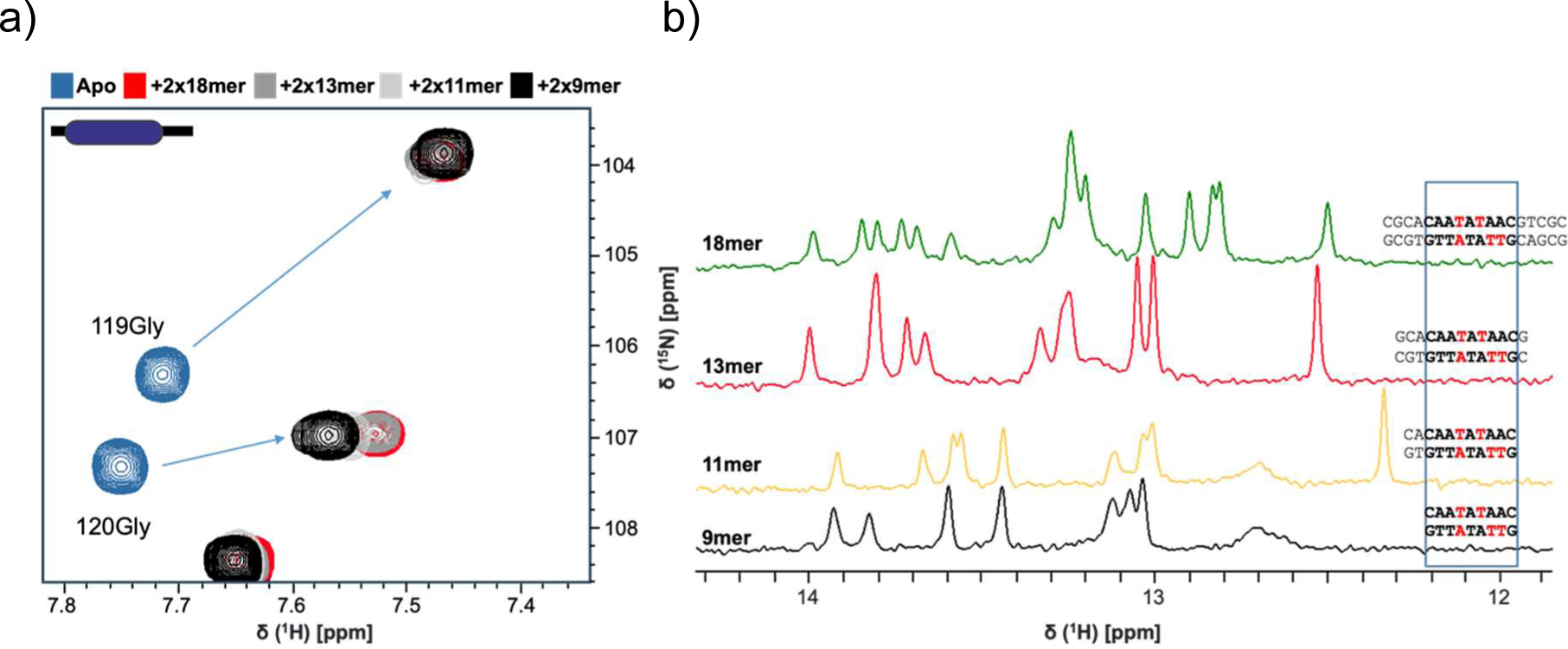
Finding the minimal dsDNA length for ARID domain binding. **a)** ^1^H-^15^N HSQC spectra of various dsDNA lengths in 2-fold excess to ARID_37-183_ (color code indicated above). **b)** Imino-proton spectra of dsDNAs used in a) show the integrity of duplexed DNA species. Sequences are given, the central AT-rich motif is boxed.

**Supplementary Fig. 4:**
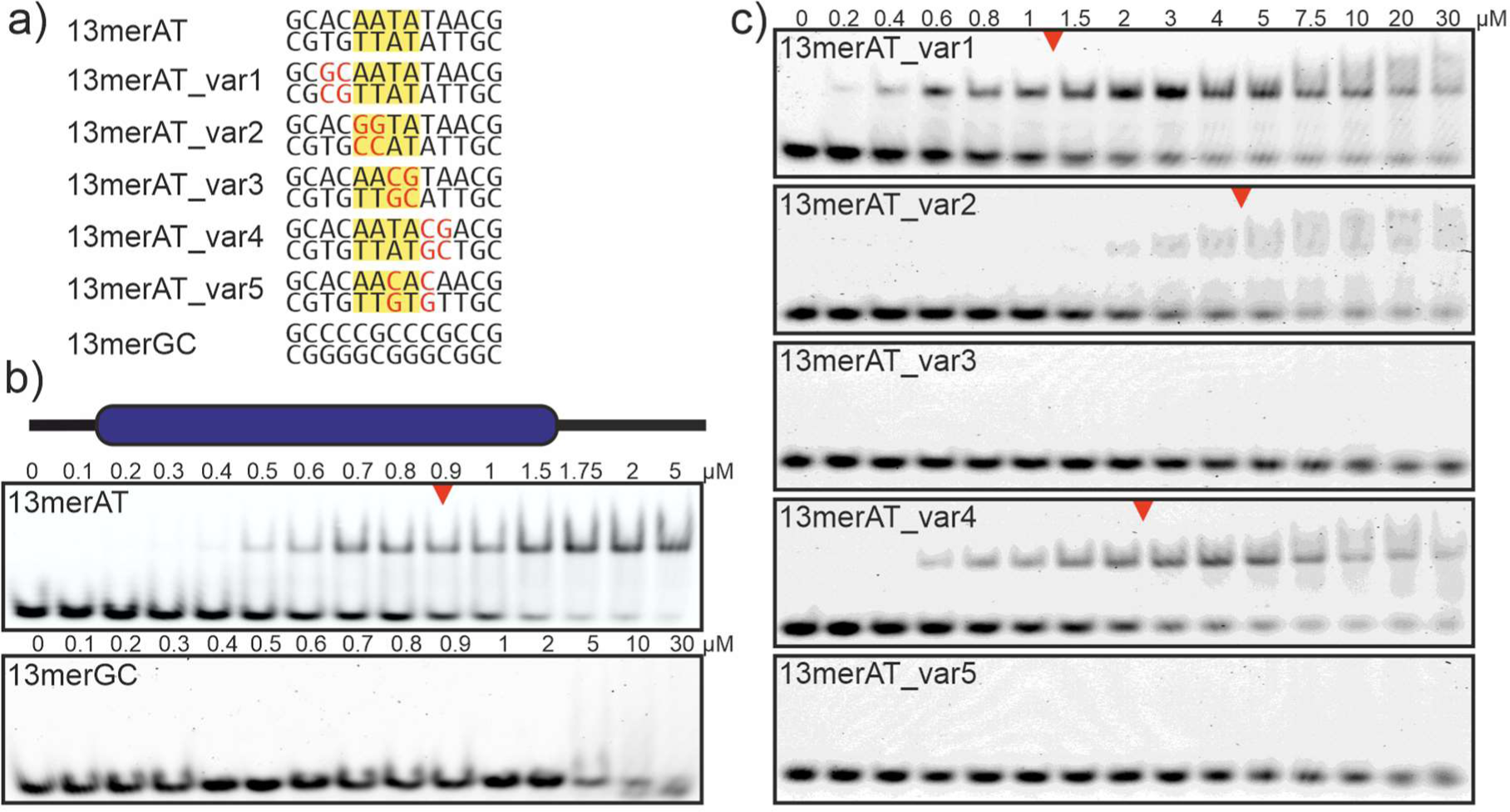
Arid5a ARID domain prefers AT-rich DNA. **a)** Sequences of DNA variants used for EMSAs. The central AT-rich motif is highlighted in yellow. Nucleotide exchanges in variants are colored red. **b)** EMSAs showing ARID_37-183_ discriminating between AT-rich and GC-rich dsDNA. EMSAs were run with 5’-fluorescently labelled dsDNA and increasing amounts of protein. **c)** EMSAs showing ARID_37-183_ discriminating between the 13merAT-variants. b) and c): Red arrows indicate approximately 50% bound DNA (as judged by optical inspection).

**Supplementary Fig. 5.**
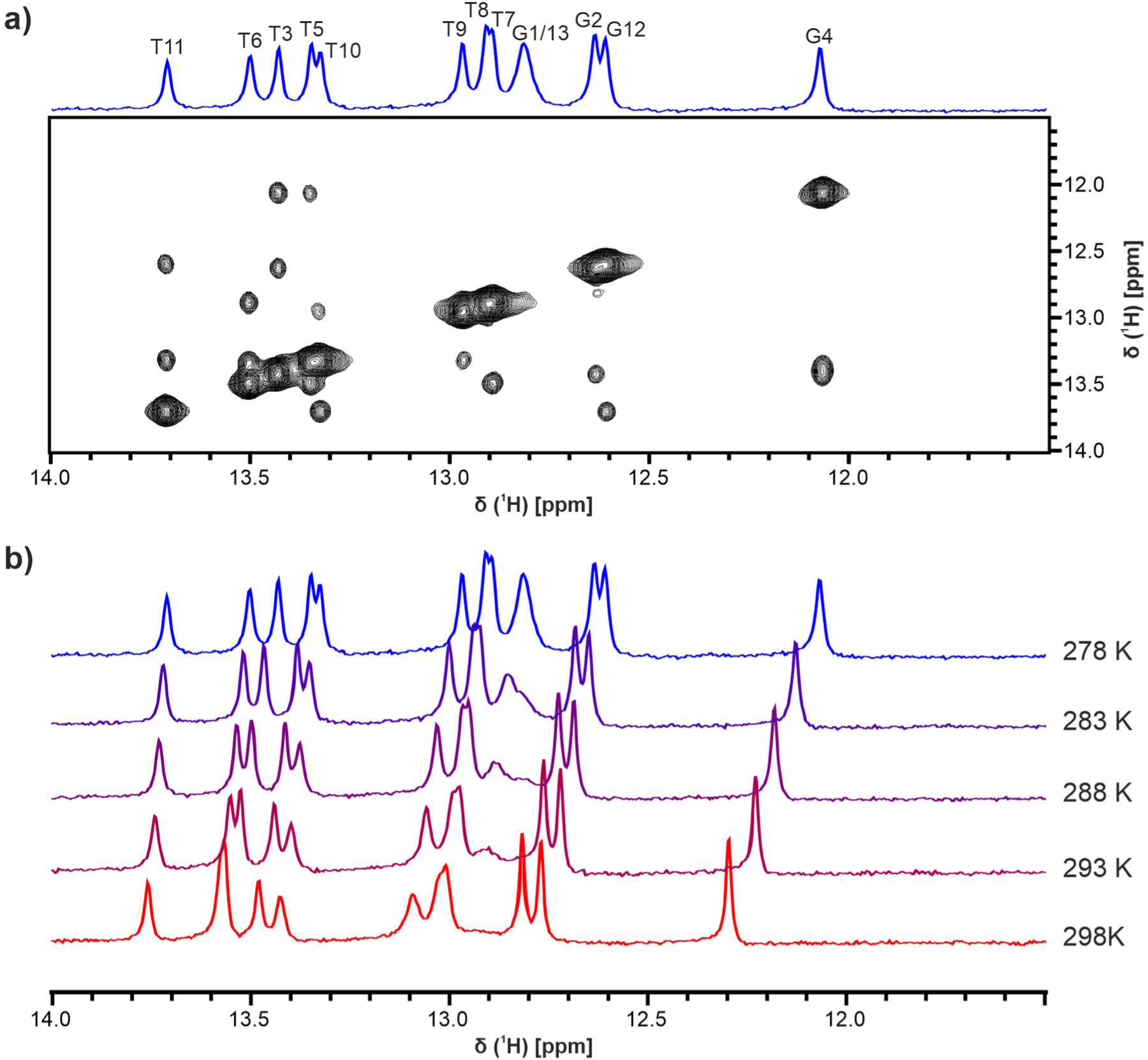
Assignment of 13merAT DNA. **a)** ^1^H-^1^H NOESY of 13merAT used to assign imino proton peaks at 278 K. **b)** Temperature series of 13merAT imino proton spectra to transfer the assignment from 278 K to 298 K (as shown in Fig. 3).

**Supplementary Fig. 6.**
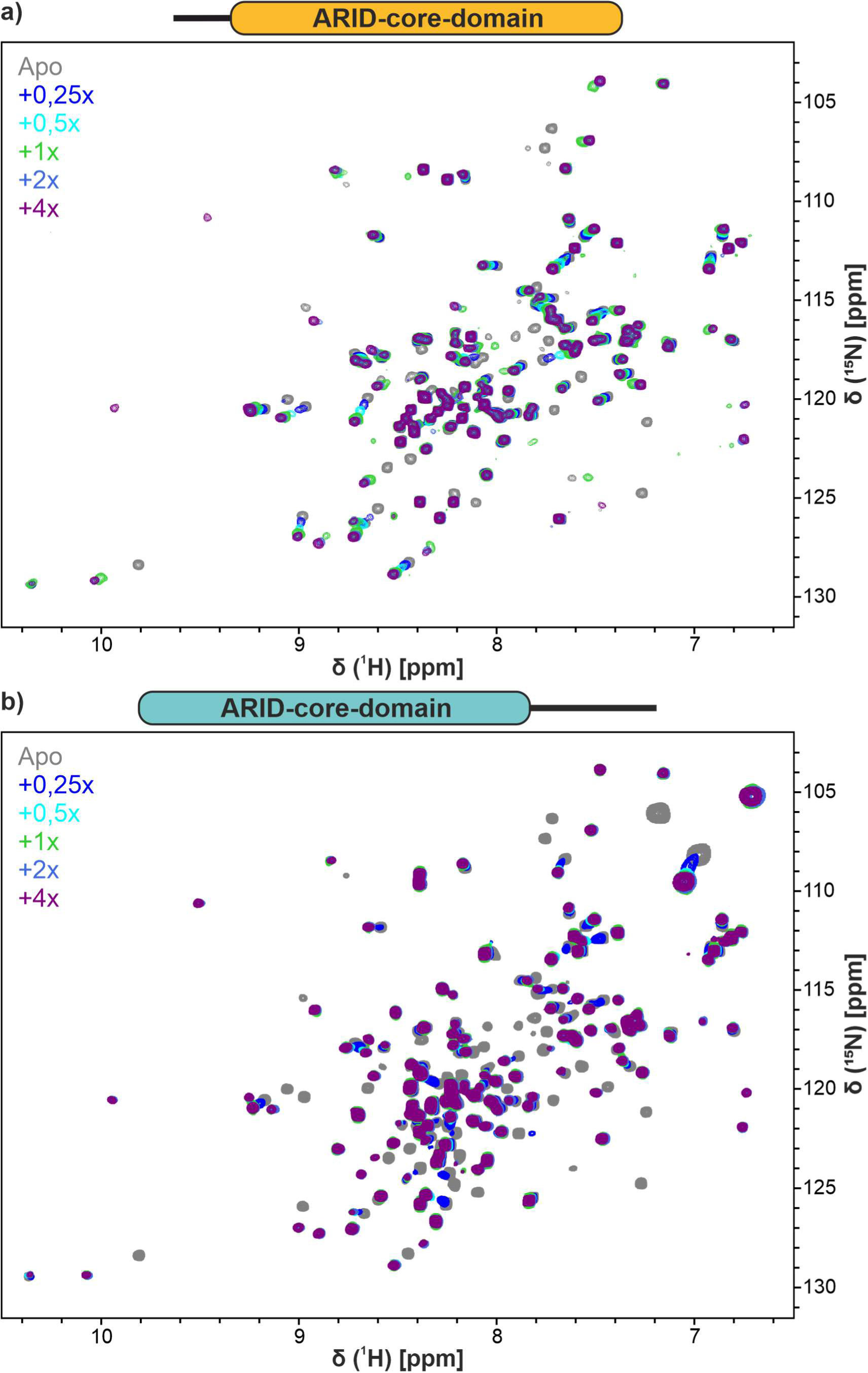
NMR-observed titrations of ARID_37-152_ and _49-183_ with 13merAT. ^1^H-^15^N HSQC spectra of ARID_37-152_ **(a)** and _49-183_ **(b)** alone (gray) and with increasing concentrations of 13merAT as indicated by the color code in upper left corner.

**Supplementary Fig. 7.**
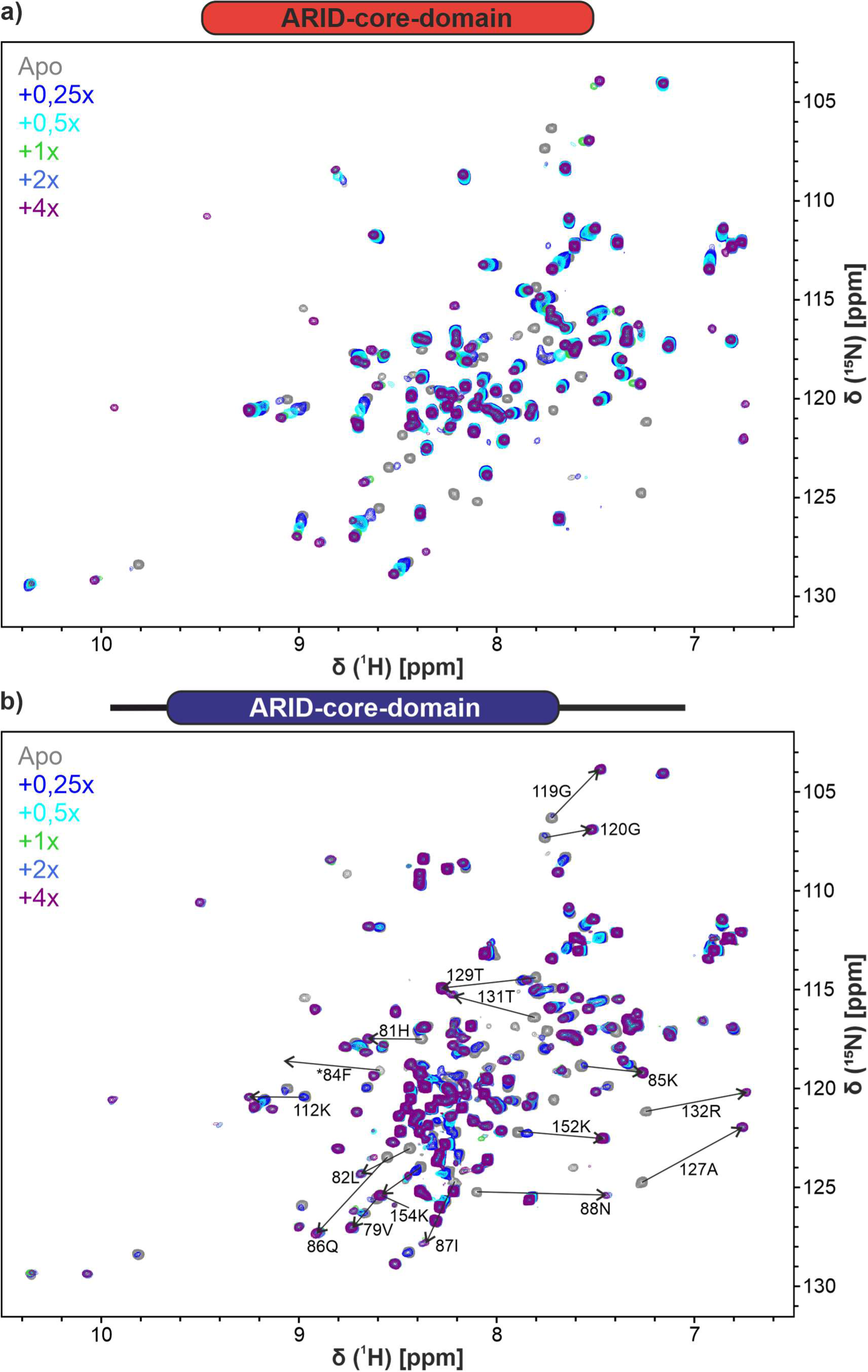
NMR-observed titrations of ARID_49-152_ and _37-183_ with 13merAT. ^1^H-^15^N HSQC spectra of ARID_49-152_ **(a)** and _37-183_ **(b)** alone (gray) and with increasing concentrations of 13merAT as indicated by the color code in upper left corner. Assignments for CSPs with values above mean+1 standard deviation (SD, according to Fig. 3b) are shown with respective trajectories indicated by arrows.

**Supplementary Fig. 8.**
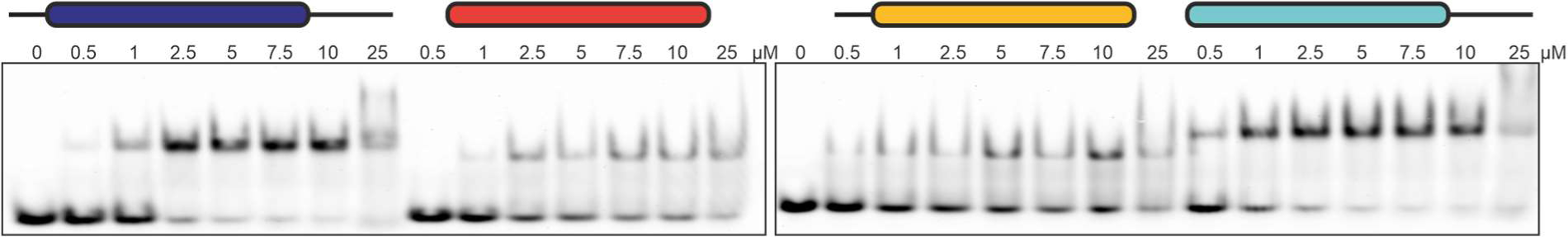
EMSAs confirm the relative affinities of core and extended Arid5a ARID constructs with 13merAT dsDNA as observed from HSQCs (Fig. 3d and Suppl. Fig. 6 and 7).

**Supplementary Fig. 9.**
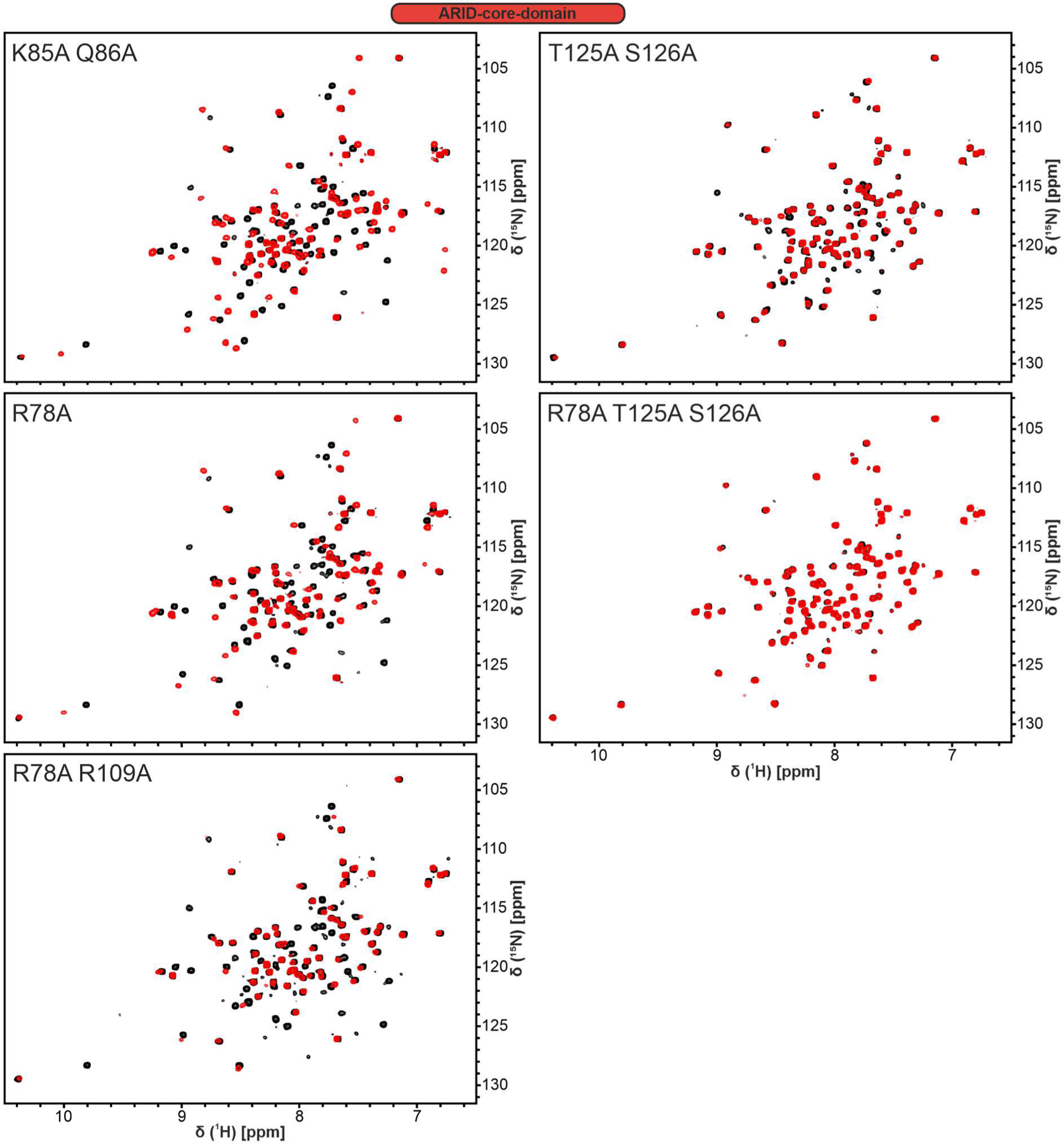
Effects of ARID_49-152_ mutants on binding to 13merAT DNA. Overlays of ^1^H- ^15^N HSQC spectra showing ARID_49-152_ mutants as indicated in the upper left corner without (black) and with 2-fold 13merAT dsDNA (red).

**Supplementary Fig. 10.**
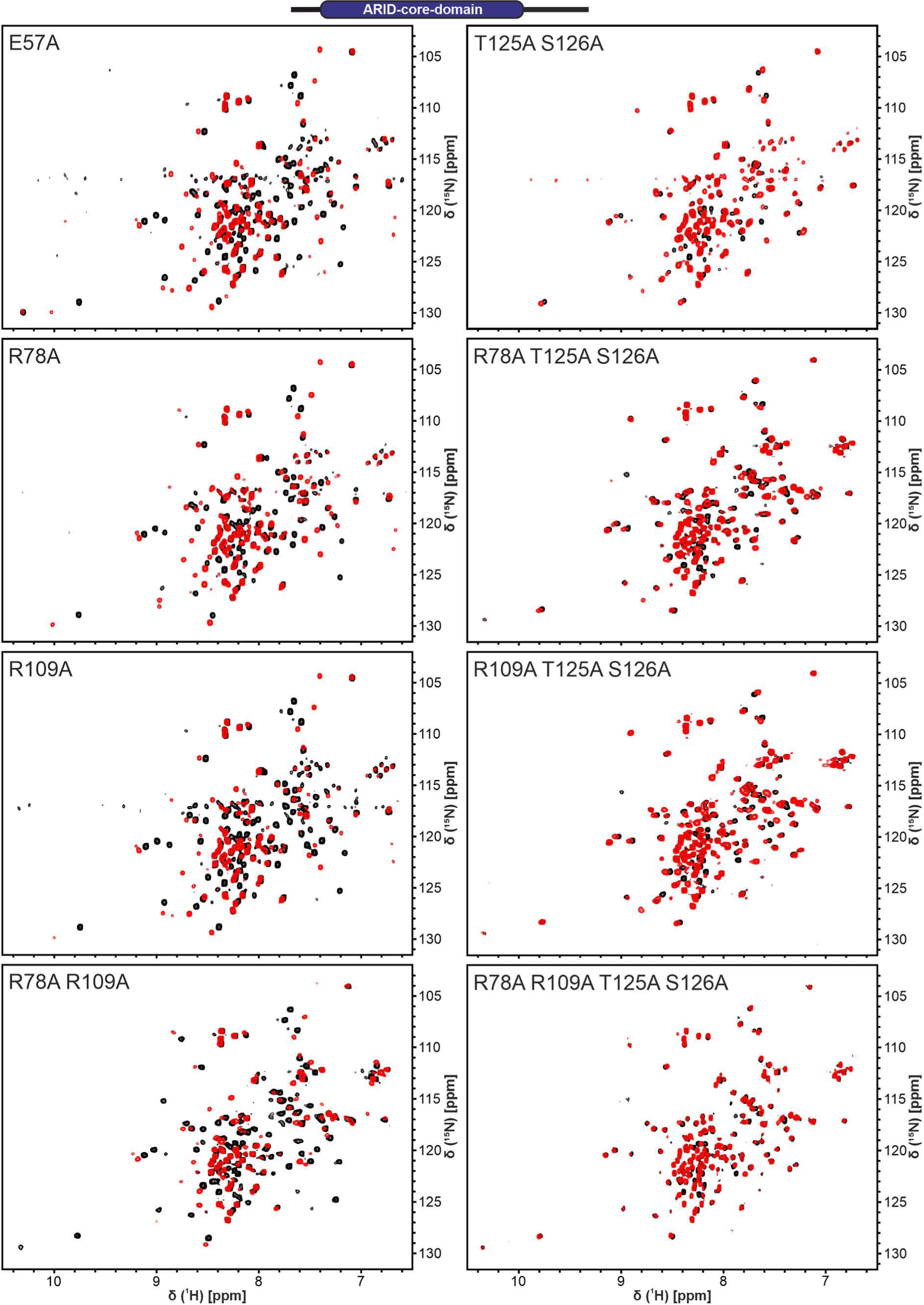
Effects of ARID_37-183_ mutants on binding to 13merAT DNA. Overlays of ^1^H- ^15^N HSQC spectra showing ARID_49-152_ mutants as indicated in the upper left corner without (black) and with 2-fold 13merAT dsDNA (red).

**Supplementary Fig. 11.**
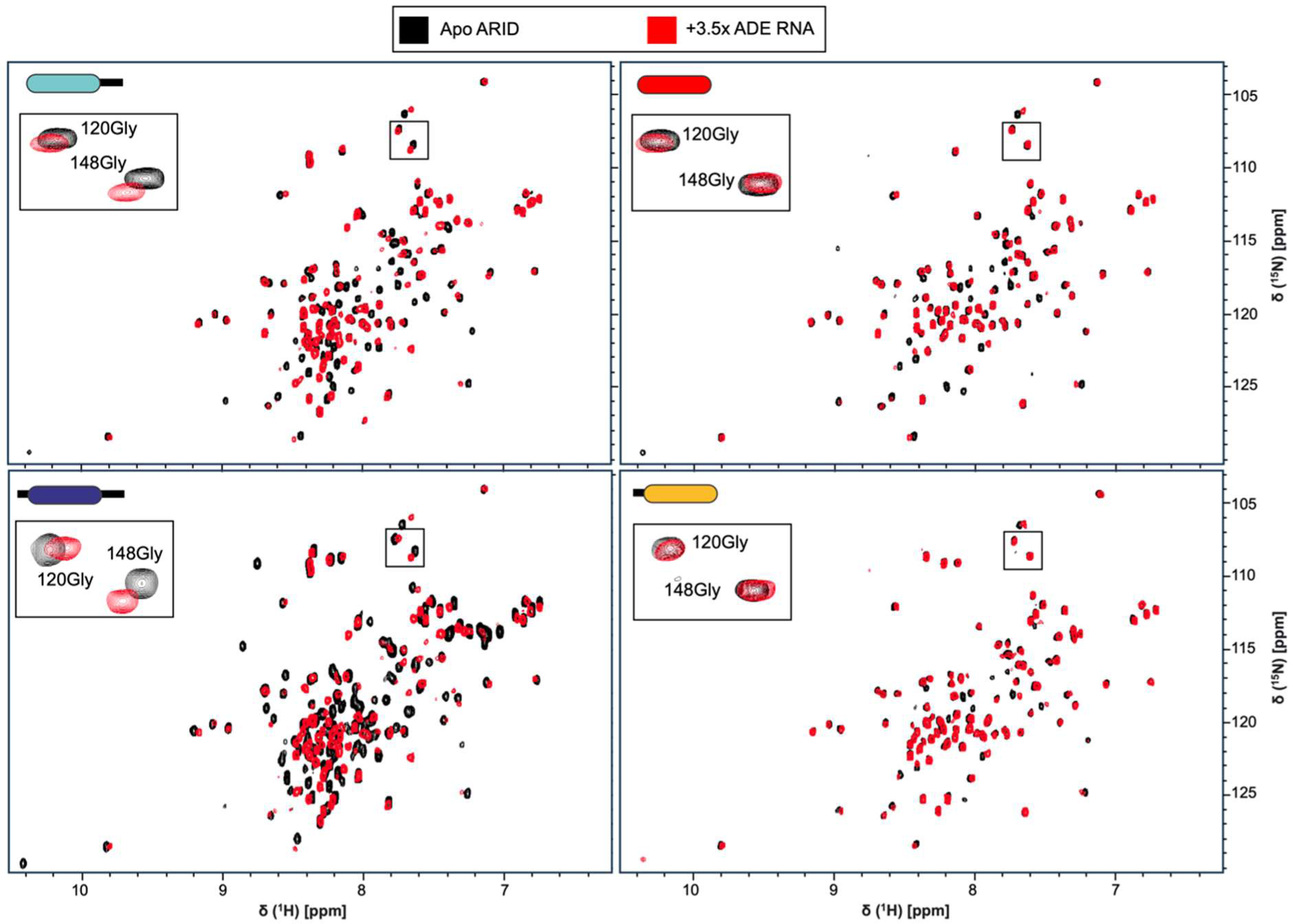
The Arid5a ARID domain binds to the *Ox40* ADE with moderate affinity but supported by the C-terminal extension. ^1^H-^15^N-HSQC spectra recorded at 298 K of 40 µM apo protein (either ARID_49-183_: top left, ARID_49-152_: top right, ARID_37-183_: bottom left or ARID_37-152_: bottom left, (black) and overlaid with the respective titration points of 3.6-fold molar excess of ADE SL-RNA (red). Insets show glycine 120, located in L2, and glycine 148, located at the C-terminal end of the structured ARID core domain.

**Supplementary Fig. 12:**
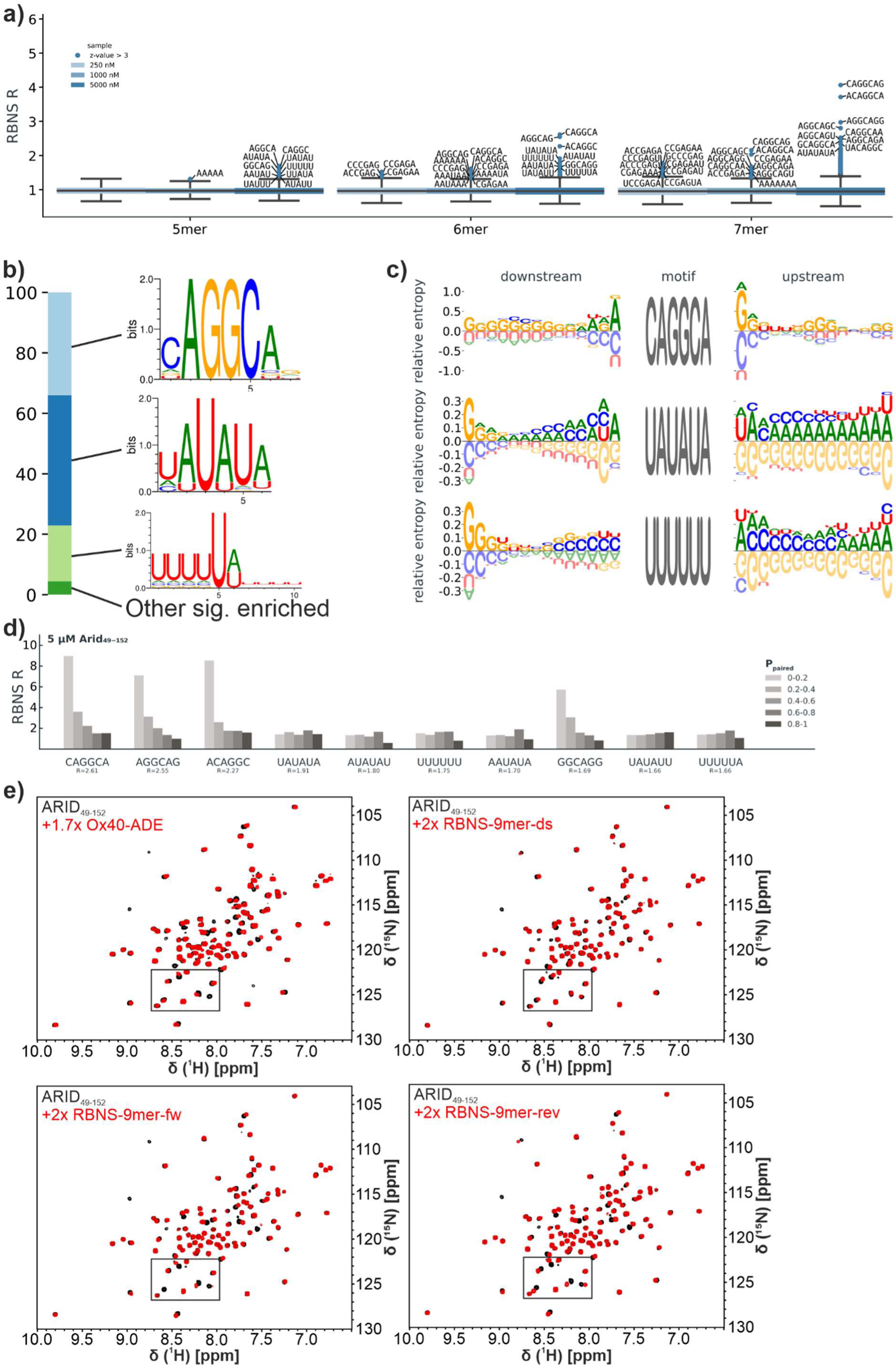
RBNS-derived sequences for ARID_49-152_. **a)** Enrichment of all different k-mers (k = 5,6,7) at 0.25, 1 and 5 µM RBP concentration. Values greater than three standard deviations above the mean are highlighted. For the highest significant ten motif sequences are given. Source data are provided. **b)** Significantly enriched sequences as logos in the most enriched 6-mers (z-score > 3). Bar plot is proportional to the summed k-mers. **c)** Sequence context: Search for complex binding motifs, which contain further recognition sequences in addition to selected enriched 6-mers at 5 µM ARID_49-152_ concentration. **d)** Structural features of the ten most enriched 6-mer motifs at 5 µM ARID_49-152_: The average P_paired_ value across the bases of the indicated motif was calculated for each occurrence in the library using RNAfold^2^. Motifs are sorted into five bins according to structure probability. The R value is calculated for each bin as the frequency in the pulldown library divided by that in the input library. **e)** Full view of ^1^H-^15^N-HSQC spectra of apo ARID_49-152_ overlaid with 1.7-fold molar excess of ADE RNA or 2-fold molar excess of RBNS-9mer RNAs for which zoom-ins are shown in Fig. 5d.

**Supplementary Fig. 13.**
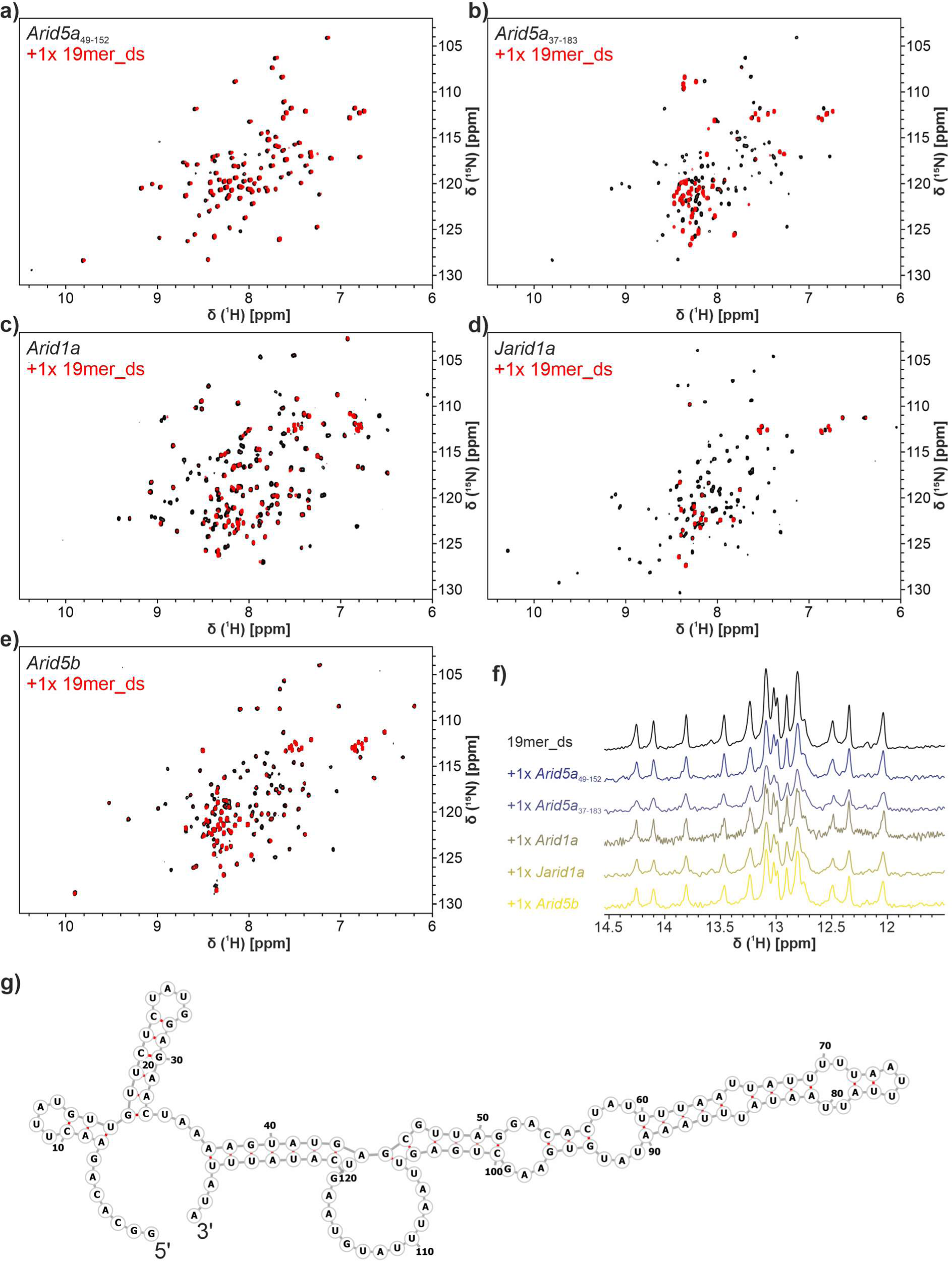
Comparative binding of ARID domains to RNA. **a-e)** Overlay of ^1^H-^15^N-HSQCs of Arid5a_49-152_ (a) or Arid5a_37-183_ (b), Arid1a ARID (c) Jarid1a ARID (d) and Arid5b ARID (e) without RNA (black) and with 1x 19mer_ds RNA (red). All protein concentrations were 50 µM. **f)** Comparison of imino proton spectra showing 19mer_ds DNA alone (top, black) or after addition of equimolar amounts of protein as indicated (black to yellow). RNA concentrations were 50 µM for samples with protein and 25 µM for the apo sample with the number of scans adjusted. **g)** Vienna RNAfold^3^ RNA secondary structure predictions of the *Il-6* 3’ UTR hub used in this study.

**Supplementary Fig. 14.**
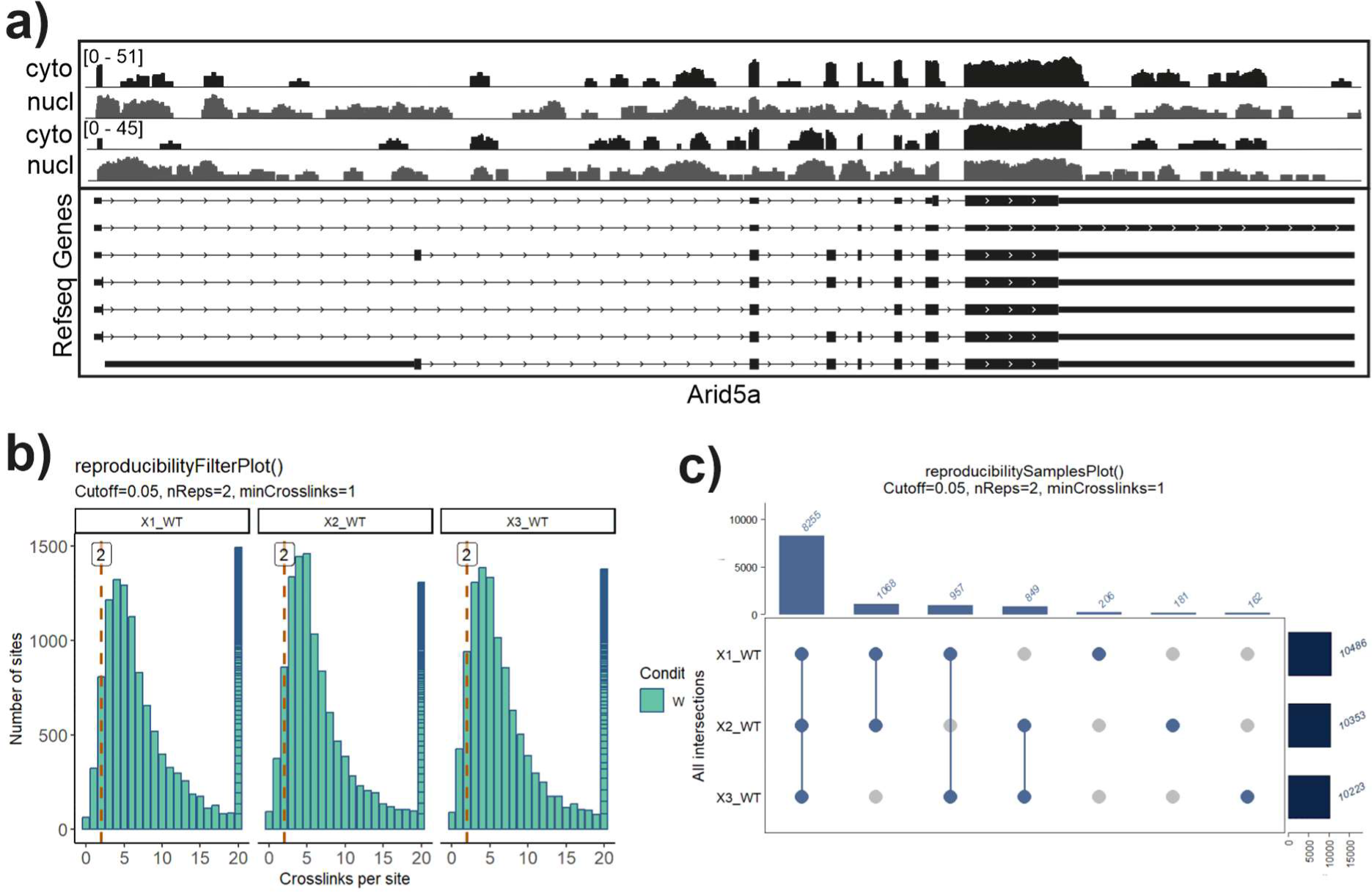
iCLIP2 analysis of Arid5a. **a)** Browser shot of nuclear and cytoplasmic RNAs (unpublished RNA-seq data) shows that Arid5a is expressed in murine P19 cells. **b)** Histogram shows the distribution of crosslink events per binding sites, indicating the minimum number of crosslink events that were required for a binding site to be called as present in a replicate. **c)** The vast majority of binding sites is shared by at least 2 of 3 replicates. Note that intergenic binding sites were removed later.

**Supplementary Fig. 15.**
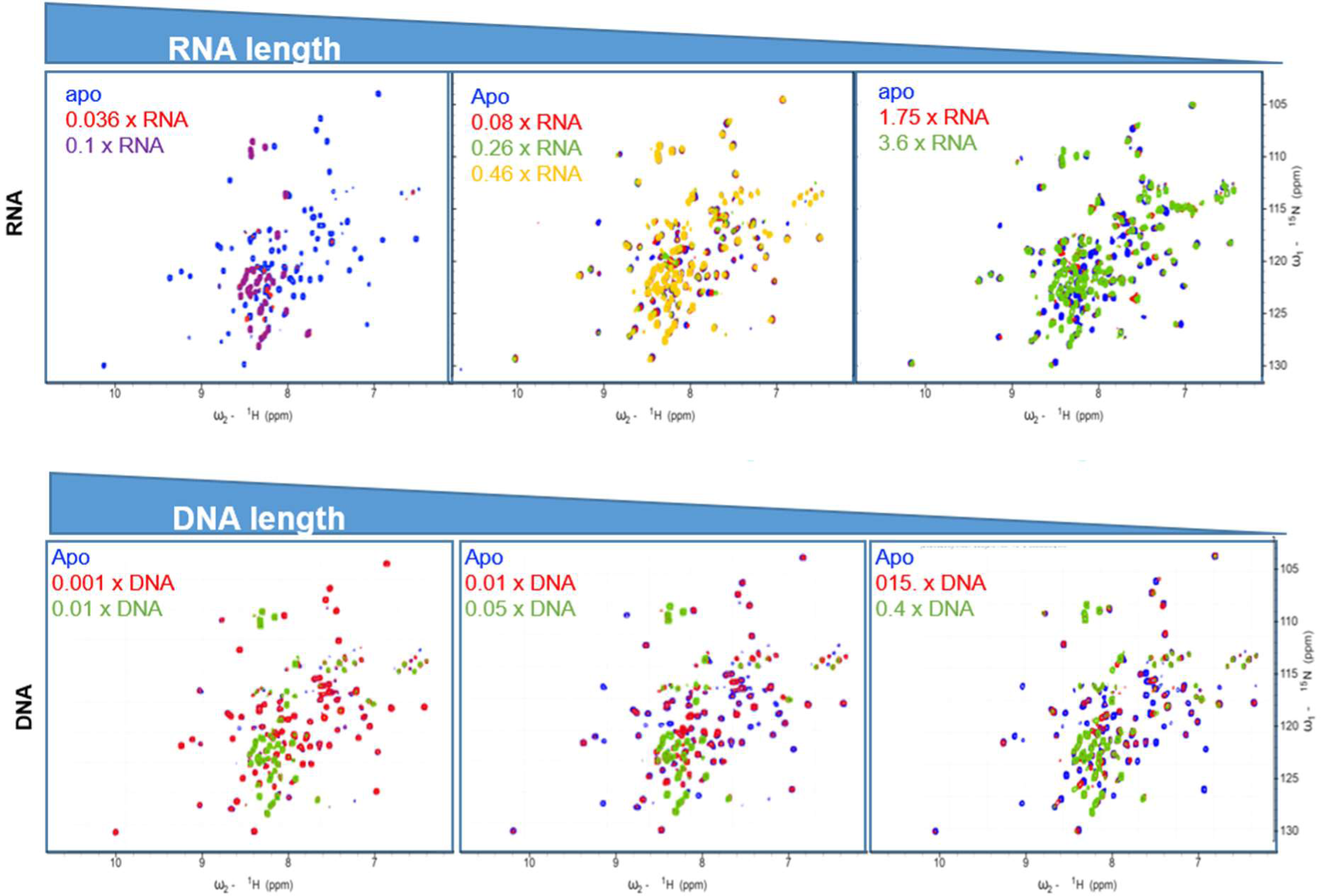
Influence of nucleic acid-length on binding to ARID_37-183_. Apparent affinities – as estimated by the degree of line-broadening in main text Fig. 7 – scale with the length of nucleic acids (NAs). Molar ratios of NA equivalents are given within the spectra. Sizes of RNAs from left to right are: 129 nt (Il_6), 60 nt (Il_6_60), 19 nt (ADE); sizes of dsDNA from left to right are: 95 nt, 70 nt, 21 nt.

**Supplementary Fig. 16.**
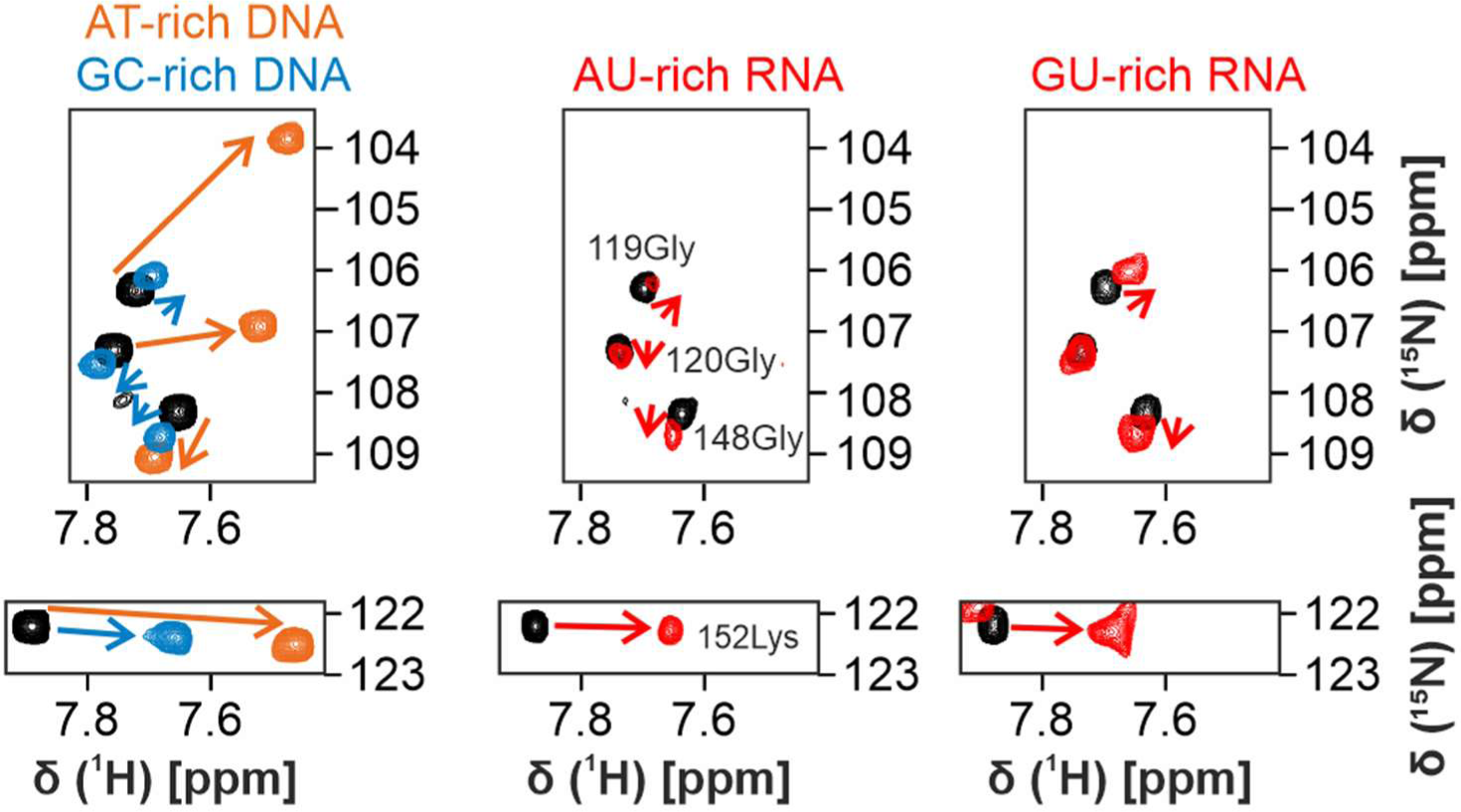
Binding of AU/GU-rich RNA by ARID_37-183_ reveals CSPs identical with non-specific GC-rich DNA binding. Shown is the direct comparison of respective insets from **Fig. 3a, Fig. 6b and Suppl. Fig. 11**. ^1^H-^15^N-HSQCs of ARID_37-183_ with 4x 13merAT (orange) and 13merGC (blue) DNAs as well as 1x 19mer_ds RNA (AU-rich, red) and 3.6x Ox40 ADE (GU-rich, red) are overlaid with the apo protein (black), respectively.

